# Host-specific platelet-activating factor acetylhydrolase selectively remodels diacylglycerophospholipids to control schistosome development

**DOI:** 10.1101/2025.04.16.649075

**Authors:** J Ertl, UF Prodjinotho, Z Rao, Anisuzzaman, J Schluckebier, Y Hamway, P Baar, M Haslbeck, CG Grevelding, S Haeberlein, FH Falcone, S Schulz, A Koeberle, C Prazeres da Costa

**Author notes:** Corresponding author: Prof. Dr. Clarissa Prazeres da Costa, Institute for Medical Microbiology, Immunology and Hygiene, Technical University of Munich Trogerstraße 30, 81675 Munich, Germany. These authors contributed equally to this work and share 1^st^ authorship (J Ertl, UF Prodjinotho).

## Abstract

**Background:** Schistosomiasis, a major neglected tropical disease, is currently treated with only one drug, praziquantel (PZQ), effective only against adult worms. However, high reinfection rates and potential development of drug resistance following widespread use of PZQ emphasize the need for a deeper molecular understanding of the host-parasite crosstalk as a basis for urgently needed novel drugs. Here, we identify and characterize a soluble host-derived schistosomicidal phospholipase that influences the diacylglycerophospholipid metabolism and, thereby, the survival and development of *Schistosoma mansoni*.

**Methods:** Large-scale proteomic screening identified host platelet-activating factor acetylhydrolase (PAFAH) as a potential schistosomicidal factor. Ultra-performance tandem mass spectrometry and electron microscopy imaging were employed to demonstrate the relevance and the lethal effects of recombinant expressed mouse PAFAH (MsPAFAH) on all juvenile stages and adult parasites from a mouse model of schistosomiasis. Quantitative lipidomic analysis revealed MsPAFAH-impaired glycerophospholipid distribution and metabolism and following free fatty acid supplementation.

**Results:** MsPAFAH was upregulated in schistosome-infected mice and exhibited potent schistosomicidal activity against all parasite life stages *ex vivo*. In contrast, human PAFAH had no effect on parasite viability. MsPAFAH treatment led to profound impairments in worm fecundity, pairing stability, reproductive organ integrity, and stem cell development. This activity was associated with substantial sex-dependent changes in ether-phospholipid composition and distribution within the schistosome tegument. MsPAFAH specifically decreased the availability of phospholipid species containing unsaturated fatty acids, namely eicosenoic (20:1) and docosatetraenoic acid (22:4), while increasing levels of respective hydrolysis (lyso) products of diacyl- and ether-phospholipids (carrying 20:1 or stearic acid (18:0)), predominantly in males. Supplementation of metabolized fatty acids C20:1/eicosenoic acid and 22:4/adrenic acid rescued the viability of female worms, confirming the essential role of the metabolism of these diacylglycerophospholipids in schistosome survival.

**Conclusions:** These findings unravel how a host phospholipase interferes with schistosome biology and development by regulating diacylglycerophospholipid availability and can thus open new avenues for schistosomiasis drug development and control.

## Introduction

Schistosomiasis, one of the 20 neglected tropical diseases (NTD), is caused by parasitic trematodes of the genus *Schistosoma* and is a significant global health challenge, affecting over 210 million people in tropical and subtropical regions worldwide (1). Despite concerted efforts to control schistosomiasis, it continues to exert a considerable burden on affected populations, particularly in resource-limited settings in sub-Saharan Africa, where it is mostly endemic. However, the limitations of PZQ, including its ineffectiveness against early larval stages and the potential emergence of drug resistance, underscore the urgent need for alternative therapeutic approaches for the populations at risk.

Schistosomiasis affects tropical and subtropical areas, and an estimated 800 million, mostly children, are at risk (1). The global burden of schistosomiasis is attributed to 3.31 million disability-adjusted life years (DALYs) per year (2). Thousands of deaths occur each year (3), and additionally, several hundred million people are struggling with residual post-treatment morbidity. In the regions with typical transmission patterns, 60 - 80% of school-age children and 20 - 40% of adults can remain actively infected despite mass drug administration (MDA) campaigns (4). Although autochthonous schistosomiasis in non-endemic regions, such as Europe, is unexpected, recent studies reported transmission in Southern Europe, indicating the probability of schistosomiasis also establishing in more moderate climate zones (5, 6). Given the vast use of a single drug only, there is concern about the emergence of drug resistance, which has been demonstrated experimentally and in a limited number of field samples (7, 8). Indeed, the WHO, in its 2030 roadmap to fight NTDs, has specifically outlined this concern to better understand the schistosome biology within the host and motivate the development of new drugs (9).

For over 30 years, in-depth investigations of mechanisms of host-parasite interaction and underlying immunopathologies *in vivo* have mainly relied on the *Schistosoma mansoni* mouse model (10). In this model, however, only about 30% of penetrated cercariae mature into fecund adult worms as mice are not the definite host for *S. mansoni* and differences in host susceptibility have been observed (10–12). We previously revealed that this loss of cercariae was not mediated by adaptive immune cells or by major factors of the complement system, like C3, C4, or C1q, or antibodies but discovered an unexpectedly strong schistomicidal effect of mouse serum on all developmental parasite stages *in vitro* (10). From the penetration of cercariae, schistosomes continuously stay in contact and bathe in the mammalian blood. Thus, it is pertinent that soluble serum factors have an obvious role in parasite development and could contribute to the host specificity of this blood-dwelling fluke (10, 11). However, most investigations so far have targeted parasite immunomodulatory and proteolytic molecules and enzymes, critical in facilitating host invasion, nutrient uptake, hatching, immune system evasion, and host physiology modulation (11, 13). In contrast, only very few host-derived proteins and their interactions with parasites have been functionally characterized (11, 14).

Among the prominent factors in host serum, enzymes play essential roles in regulating energy levels. By catalyzing the hydrolysis and modification of key molecules to provide nutrients, generating bioactive mediators, and influencing cell metabolism, host enzymes, e.g. human lipoxygenases and peroxiredoxins, are central to the survival of parasites (11, 15–17). As such, host enzymes are, in general, essential targets for regulating cell metabolism in various research areas (18). In cancer research, targeting glucose and lipid metabolic enzymes has emerged as a promising strategy for developing antineoplastic drugs (19, 20). In helminths, especially for schistosomes, lipid metabolism plays a crucial role in the survival and pathogenesis of the worm, making it a relevant target for novel therapeutic strategies (11, 21, 22). Schistosomes rely on lipid metabolism to acquire essential nutrients for their growth, development, and reproduction, e.g. for, egg production and maturation (23–25). Essentially, lipids and phospholipids (PL) are important components of the schistosome membrane, contributing to its structure, integrity, functions, and host immune evasion (22, 26). Furthermore, schistosomes are unable to synthesize fatty acids and sterols *de novo* (25, 26). Hence, disruption of schistosome PL metabolism can impair nutrient uptake, ion transport, and communication with the host environment, ultimately leading to parasite death (15, 25). Thus, host phospholipases, a group of enzymes capable of hydrolyzing PL, are particularly interesting as they potentially represent anti-schistosomal molecules (11, 23).

In this study, through multifaceted molecular approaches and proteomic analyses, we unmasked that the lipoprotein-associated phospholipase A_2_, platelet-activating factor-acetylhydrolase (PAFAH) of host origin, is one of the major serum factors, which critically influences the survival of mammalian host-dwelling stages of schistosomes. We demonstrated *ex vivo* that recombinant mouse, but not human, PAFAH, disrupts tegument integrity and essential parasite structures, interferes with lipid metabolism and homeostasis, and impairs key physiological processes crucial for parasite survival and development. These findings shed light on mechanisms underlying host preference and specificity of schistosomes and enable further research into the identification of the mode of action of PAFAH. This may guide the development of novel drugs, such as small molecules interfering specifically with the PL metabolism of the parasite.

## Methods

### Ethical statements, animals, blood samples, and serum preparation

The animal and mouse serum investigations were approved by the Bezirksregierung Oberbayern (license number AZ 55.2-1-54-2532-145-17). NMRI mice were purchased (Envigo, Germany) or bred in-house. Animals were maintained according to national and EU guidelines 86/809 under specific pathogen-free conditions at the Institute for Medical Microbiology, Immunology, and Hygiene (MIH) Animal facility. Animals (6-8 weeks of age) of both sexes were used. Sera from infected and non-infected mice were prepared from blood collected by venipuncture in non-medicated Falcon tubes. The blood was centrifuged for five minutes, and serum was collected separately or pooled when indicated and stored at -20°C until use.

Investigations using human serum were approved by the local ethical committee of the Technical University of Munich (TUM) (Reference: 215/18S), and all individuals, included in the study, consented enrollment. Human serum was prepared from the blood of healthy female and male volunteers with no previous history of schistosomiasis. Fresh blood was clotted at room temperature for 30 min and centrifuged at 1,845 x g for 20 min. Serum was collected, pooled, and stored at -20°C until further use.

### NTS generation, culture, and advanced larval stage development

Newly Transformed Schistosomula (NTS) were generated as previously described (27, 28). For *in vitro* culture, approximately 100 NTS in 150 µl HybridoMed (HM) culture medium (HybridoMed Diff 1000 (Biochrom GmbH, Germany) supplemented with 200 U/mL Penicillin, 200 μg/mL Streptomycin (Sigma-Aldrich) were plated in 96-well flat-bottom tissue culture plate and incubated at 37 °C in 5% CO_2_ atmosphere to rest for 48 hours. For long-term culture and advance stage development, 100 NTS were resuspended in HM supplemented with 20% human serum (HM+HSe) and cultured at 37 ̊C, 5% CO_2_ atmosphere for 25-30 days post-transformation (p.t.) (27). To guarantee optimal culture conditions, culture-medium was replaced twice a week. Developmental stages (Lung stage (LuS), Early liver stage (eLiS), and Late liver stage (lLiS) schistosomula were examined by bright field microscopy using an inverted Axiovert 10 microscope (Zeiss) as previously published (10, 27–29).

### Collection of *ex vivo* adult worms

3-4-week-old male NMRI mice were purchased from Envigo Germany and infected subcutaneously with approximately 200 *S*. *mansoni* cercariae (NMRI strain) per flank. After 6– 7 weeks post-infection, mice were euthanized, and worms were collected from the portal and mesenteric veins via conventional perfusion (30). Adult worms were placed in a petri dish with HM, washed once, and 10 worms/well were gently transferred into a 6-well plate with 2 ml HM supplemented with 20% HSe and rested for 24h at 37 °C, 5% CO_2_ before *in vitro* testing with sera and PAFAH.

### Recombinant protein production and purification and sequence alignment and homology modeling

The short-listed proteins identified as most likely active killing candidates (major urinary protein 10 (MUP10), fibroblast activation protein (FAP), and adiponectin) were cloned in pET28b without signal peptides using the NdeI and XhoI restriction sites and fusing the vector encoded N-terminal Histag. The recombinant proteins were expressed in *E. coli* BL21 strains and purified using metal affinity, ion exchange, hydrophobic interaction, and size exclusion chromatography. Recombinantly expressed in *E. coli* and purified mouse PAFAH (MsPAFAH) (PLA2G7, N-terminal His tagged, abx068540) and human PAFAH (HuPAFAH) (PLA2G7, N-terminal His tagged, abx068539) proteins were both purchased from Abbexa (Cambridge, UK), the purity checked, and dissolved in PBS, aliquoted at 1 mg/ml, and stored at -80 ̊C until use. Protein sequences and alignment were searched with UniProt and Clustal Omega tools, and homology modeling of both HuPAFAH and MsPAFAH structures and motif, domain, and active sites were performed using ExPASy (Swiss Bioinformatics Resources Portal) and AlphaFold2 modeling (UniProt) tools.

Structures of HuPAFAH (AA 54-427) and MsPAFAH (AA 22-426) (HUGO Gene name: PLA2G7) based on the 1.50 Å structure of HuPAFAH (3D59.pdb) and the AlphaFold2 predicted structure for MsPAFAH (available on UniProt entry Q13093) were visualized using UCSF ChimeraX 1.2.5. The N-terminal signal sequences in both proteins are not shown, as are sections not imaged in the 3D59 X-ray structure.

### *In vitro* assays with serum, PAFAH, and viability scoring

For *in vitro* culture of NTS, parasites were cultured in 96-well plates supplemented with 150 µl HM. Each well contained approximately 100 NTS. After 48h resting, wells with an average score ≥ 2.00 scoring points were selected for *in vitro* compound screening as previously established (27, 28). These were washed with HM to remove metabolites and waste generated during cultivation. Following the manufacturer’s instructions, PAFAH was dissolved in PBS and added at 1–50 μg/ml final concentrations. As a negative control, HM was adjusted to the same concentration of PBS as used for the PAFAH test. 20% mouse serum dissolved in HM (MSe) was used as a positive control to compare morphological effects between phenotypes and PAFAH. The worms were incubated at 37 ̊C and 5% CO_2_ for 168 h. PAFAH-induced morphological effects and viability scores were assessed after 24, 72, and 168 h using an inverted Axiovert 10 microscope (Zeiss, Germany) at 10x magnification. The viability of parasites in culture was scored as previously described (27, 28). Briefly, the scoring system is based on three main viability criteria: morphology, granularity, and motility, ranging from 0.00 to 3.00, with a score of 0.00 representing dead parasites and a score of 3.00 for entirely healthy, motile larvae. As the score represents an average of all NTS per well, subtle differences in viability were assessed by subdividing viability scores into 0.25 steps (e.g., 2.00, 2.25, 2.50, 2.75).

For long-term culture and advanced-stage development, 100 NTS were kept in 96-well plates and resuspended in HM supplemented with 20% human serum (HM+HSe). They were cultured at 37 ̊C, 5% CO_2_ until indicated time points post-transformation (p.t.). Developmental stages (lung stage schistosomula (LuS) and early (eLiS) and late liver schistosomula (lLiS)) were generated and characterized according to previously published morphological criteria (27, 29, 31).

For compound assessment, each well contained approximately 100 parasites with an average score ≥ 2.00 to be selected for screening (27, 28). After selection, juveniles were washed with HM to remove metabolites and waste generated during cultivation. PAFAH was dissolved in PBS and added in final concentrations of 1–50 μg/ml as indicated, and *in vitro* assays and viability scoring were performed as described above.

To inhibit PAFAH activity *in vitro* we developed an inhibition assay. For this, Varespladib (VAR) (Sigma, SML1100), a potent and selective inhibitor of both human and mouse secretory phospholipase A2 (sPLA2) (32, 33) was prepared as 10 mM stock solution in dimethyl sulfoxide (DMSO) to be used at a final concentration of 5 nM diluted in HM. For the *in vitro* assay, VAR was pre-incubated with MSe or MsPAFAH (at respective concentrations as above) for 1 h at 37°C before incubation with NTS for 72 h.

### *In vitro* assays with adult worms

As previously described, *S. mansoni* adult worms of both sexes were flushed out from infected mice (28, 34) and rested for 24h at 37 ̊C and 5% CO_2_. For the *in vitro* culture of adult couples, worms were cultured in 6-well plates supplemented with 2 ml HM for 10 worm couples or 10 females or males per well. PAFAH was dissolved in PBS and added in final 1–50 μg/ml concentrations as indicated. HM was used as a negative control with 20% MSe as a positive control. The worms were incubated at 37 ̊C and 5% CO_2_ for 72h; medium and compounds were exchanged every 24h. PAFAH-induced morphological effects were assessed every 24h using an inverted Axiovert 10 microscope (Zeiss) at 10x magnification.

Adult worm viability was scored as above and based on the recommendations by WHO-TDR (35, 36), with the scores 3 (normal viability), 2 (reduced viability), 1 (minimal viability), and 0 (no movement within 30 sec was considered dead). Viability scores were subdivided into 0.25 steps. The scoring system is based on criteria like attachment to the well ground, granularity, and movement. The pairing status was observed to assess reproductive parameters for cultured worm couples, and eggs were counted daily. After 48h-72h, worms were either processed for confocal laser scanning microscopy (CLSM), scanning electron microscopy (SEM), and HPLC analysis or subjected to RNA extraction for quantitative real-time PCR (qPCR) analysis.

For phospholipid (PL) and free fatty acid (FFA) supplementation assays, *S. mansoni* adult worms were flushed out from infected mice and separated by sex, and 5 worms/well were plated as above in 2 ml DMEM (Gibco, ThermoFisher Scientific). Then, worms, treated with 15 µg/ml MsPAFAH and 20% MSe, were supplemented in parallel with 50 µM of C20:1/eicosenoic acid or 22:4/adrenic acid (Merck KGaA, Germany) to compensate the depletion of PC(16:0_20:1) and PE(18:0_22:4), or 18:0/stearic acid or 14:0 DMPC (1,2-dimyristoyl phosphatidylcholine) (Merck KGaA, Germany) as controls for 72h and viability was assessed as described above.

### Large-scale fractionation of mouse serum

Sera were harvested from naive mice derived from our in-house breeding facility or Envigo and diluted with PBS before fractionation using the Superdex 200 column (GE Healthcare Life Sciences). Collected fractions were pooled (F1, F2, F3, F4, and F5) according to dominant chromatographic peaks, and the volume was adjusted to the initial fractionated mouse serum volume and used at 20% for the NTS *in vitro* assay as described above.

### Mass spectrometry of active fractions and identification of active components

For the identification of proteins present in the active fractions of mouse serum, the respective fractions were separated by SDS-PAGE. The resulting lanes of the SDS-PAGE were divided into 3-5 parts and sliced out of the gel. The proteins in the gel slices were reduced and alkylated before in-gel tryptic digest (using MS-grade trypsin, Promega, V511). The resulting peptides were extracted from the gel and applied to an UltiMate 3000 nano HPLC System, loading the peptide solution onto an Acclaim PepMap RSLC C18 trap column (Trap Column, NanoViper, 75 μm x 20mm, C18, 3 μm, 100 Å, Thermo Scientific) and separating the peptides on a PepMap RSLC C18 column (CoIP1: 75 μm x 50 mm C18, 2 μm, 100 Å, Thermo Scientific). Linear gradients from 4 % (vol/vol) to 35 % (vol/vol) acetonitrile with 0.1 % formic acid were applied to elute the peptides, which were subsequently analyzed in an Orbitrap Q Exactive plus (Thermo Scientific) mass spectrometer. Full scans of collision-induced dissociation MS^2^ (scan 2 mode) of the most intense ions were recorded throughout the elution gradient.

The mass spectrometry data from each sample were searched against the Swiss-Prot mouse database downloaded from UniProt using the Sequest HT Algorithm implemented into the “Proteome Discoverer 1.4” software (Thermo Scientific). The search was limited to peptides containing a maximum of two missed cleavage sites and a peptide tolerance of 10 ppm (parts per million) for precursors and 0.04 Da for fragment masses. Proteins were identified with at least one unique peptide with a target discovery score of higher than one. For further evaluation, three independent datasets resulting from biological replicates were compared.

For candidate selection for bioassay, the identified proteins were screened for the following selective criteria using Gene Ontology: 1) type and state (secreted), 2) function and activity (protein-, lipid-, phospholipid-binding activity, calcium-dependent), 3) linked to protein- or lipid-metabolism.

### Confocal laser scanning microscopy (CLSM)

For morphological analysis by CLSM, worms were fixed and stained for 30 min with carmine red (CertistainH; Merck, Germany) as described before (37, 38). After treatment, the worms were fixed in AFA (95% alcohol, 3% formalin, and 2% glacial acetic acid) and stained with 2.5% hydrochloric carmine (Certistain, Merck). The specimens were preserved as whole mounts in Canada balsam (Merck) on glass slides. Stained worms were examined on an inverse CLSM (Confocal Olympus FV3000). Carmine red was excited with an argon-ion laser at 488 nm. Laser power and gain and offset of all photomultiplier tubes (PMTs) were optimized to minimize possible bleaching effects and complete range intensity coding using the CLUT function (color look-up table) of the Olympus FV3000 software. Background signals and optical section thickness were defined by setting the pinhole size to airy unit 1.

### Scanning electron microscopy (SEM)

After culture, parasites were collected in protein low-binding Eppendorf tubes, fixed in 4% glutaraldehyde / 0.2 M cacodylate, and washed with 0.4 M saccharose / 0.2 M cacodylate. Following fixation, worms were treated with 1% osmic acid / 0.1 M cacodylate and washed for a second time in distilled water, followed by ethanol dehydration at 30%, 50%, and 70% concentration. The next day, dehydration in 80%, 90%, 96%, 99.8%, and 100% ethanol continued. After all, samples were air-dried in a CPD030 critical point dryer (BAL-TEC AG; Balzers, Liechtenstein), mounted, and coated with gold sputter before imaging with a Gemini DSM 982 (Carl Zeiss Microscopy; Oberkochen, Germany), operating at 3 kV.

### MALDI-MS Imaging

The sectioning protocol for the samples was adapted from Mokosch et al. (39). Instead of 15 μL, 30 μL of gelatine solution were used. The sections of the male samples had a thickness of 20 μm, and the female sections were 16 μm thick. The samples were measured with an atmospheric-pressure MALDI imaging ion source (AP-SMALDI5 AF, TransMIT GmbH) coupled to an orbital trapping mass spectrometer (Q Exactive HF, Thermo Fisher Scientific). Measurements were performed in positive ion mode using a mass range of m/z 250-1000, a mass resolution of R = 240,000 @ m/z 200, and a spatial resolution (i.e., step size) of 7 µm. Internal mass calibration was achieved using the lock-mass feature of the orbital trapping mass spectrometer, resulting in a mass accuracy of < 3 ppm. For data analysis, raw data of all measurements were stitched together and converted to imzML imaging data file format. Annotation of lipid signals ([M+H]^+^, [M+Na]^+^, [M+K]^+^) was performed by Metaspace, a fully automated metabolite annotation platform for imaging data (40), using the databases of HMDB-v4, LipidMaps-2017-12-12 and SwissLipids-2018-02-02 and a ± 3 ppm m/z window.

For semi-quantification, regions of interest (ROI) comprising the tissue of each worm were defined. Total ion count (TIC) normalized images of annotated lipids (+/-3 ppm) containing the ROIs of all samples were exported from MIRION software in CSV-Format (41). In Matlab, the intensity sum of each lipid signal was calculated per ROI and divided by the ROI pixel number for each sample.

### Extraction and quantitative analysis of phospholipids

Phospholipids were extracted from cell pellets by the sequential addition of PBS pH 7.4, methanol (spiked with internal standards), chloroform, and saline (final ratio 14:34:35:17). The organic layer was evaporated using an Eppendorf Concentrator Plus system (Hamburg, Germany; non-polar phase: high vapor pressure application mode), stored at -20°C, and dissolved in methanol before analysis (42, 43). Internal standards: 1-pentadecanoyl-2-oleoyl(d7)-sn-glycero-3-phosphocholine [PC(15:0_18:1-d7), Avanti Polar Lipids, Alabaster, AL] and 1-pentadecanoyl-2-oleoyl(d7)-sn-glycero-3-phosphoethanolamine [PE(15:0_18:1-d7), Avanti Polar Lipids].

Phospholipids were separated on an Acquity UPLC BEH C8 column (130 Å, 1.7 μm, 2.1×100 mm, Waters, Milford, MA) using an ExionLC AD UHPLC system (Sciex, Framingham, MA) and detected by a QTRAP 6500^+^ mass spectrometer (Sciex) equipped with an IonDrive Turbo V Ion Source and a TurboIonSpray probe for electrospray ionization (43). Chromatographic separation was performed at 45°C and a flow rate of 0.75 ml/min using mobile phase A (water/acetonitrile 90/10, 2 mM ammonium acetate) and mobile phase B (water/acetonitrile 5/95, 2 mM ammonium acetate). The gradient was ramped from 75 to 85% B over 5 min and then increased to 100% B within 2 min, followed by isocratic elution for another 2 min. Optimized MS source and compound parameters for PC and PE are shown in Table 1. The QTRAP 6500^+^ system was operated in negative ionization mode using scheduled multiple reaction monitoring (MRM). The transitions from [M-H]^−^ to both fatty acid anions were monitored (42, 43) to detect PE and PC species. The instruments were operated by Analyst 1.7.1 (Sciex), and the mass spectra were processed by Analyst 1.6.3 (Sciex).

**Table 1:** Source and compound parameters.

| Parameters | PC (diacyl) | PE (diacyl- and ether-) |
| --- | --- | --- |
| Curtain gas (CUR) | 40 psi | 40 psi |
| Collision gas (CAD) | medium | medium |
| Ion spray voltage (IS) | -4,500 V | -4,500 V |
| Heated capillary temperature (TEM) | 350°C | 650°C |
| Sheath gas pressure (GS1) | 55 psi | 55 psi |
| Auxiliary gas (GS2) | 75 psi | 75 psi |
| Declustering potential (DP) | -44 V | -50 V |
| Entrance potential (EP) | -10 V | -10 V |
| Collision energy (CE) | -46 eV | -38 eV |
| Collision cell exit potential (CXP) | -11 V | -12 V |

Combined peak areas from the transitions to the two fatty acid anions were calculated for the quantitative analysis of PE and PC. To calculate absolute lipid quantities, signals were normalized to a subgroup-specific internal standard [PE(15:0_18:1-d7) or PC(15:0_18:1-d7)]. Relative intensities of phospholipids are given as a percentage of all species detected in the corresponding phospholipid subclass (= 100%). Cellular proportions of phospholipid subgroups were calculated by summarizing the relative intensities of individual species.

### ELISA and PAFAH protein activity assay

According to the manufacturer’s instructions, PAFAH concentrations in infected and non-infected mouse serum were determined using commercially available enzyme immunoassay kits (Abbexa). PAFAH enzyme activity in different sera and of the recombinantly expressed protein MsPAFAH (44 µg/ml) was assessed using the PAF Acetylhydrolase Assay Kit (Cayman Chemical) according to the instructions provided by the manufacturer and with HuPAFAH as positive control.

### RNA isolation and RT-qPCR analysis

Freshly perfused or *in vitro*-cultured worm couples were separated by gender, and 5 worms per gender were incubated in different culture conditions in du-/triplicate. After the indicated time points, RNA was extracted from male and female worms using the Monarch^®^ Total RNA Miniprep Kit (New England Biolabs) according to the manufacturer’s protocol. Reverse RNA transcription in cDNA was performed using the Quantitect RT-Kit (QIAGEN). Expression levels of the *S*. *mansoni* orthologs of the stem cell markers *nanos-1* (Smp_055740) and *fgfr1* (Smp_157300) and apoptosis *bcl2* (Smp_072180) were determined by RT-qPCR using the KAPA SYBR® FAST qPCR Master Mix (2X) Kit (Sigma-Aldrich). All samples were pipetted in technical triplicates. Ct values were normalized against the geometric mean of the reference gene *letm1* (Leucine zipper/EF-hand-containing transmembrane protein 1) (Smp_065110), selected based on stable expression in both sexes (44). Relative expression levels were calculated by the delta-delta Ct method (45) or by expressing the data as n-fold difference by the formula: relative expression = 2^−delta Ct^ × f, with f = 1 as an arbitrary factor. The following primers were used, which were confirmed by test qPCRs to have efficacies between 0.9–1: LETM1_ forward (fw) 5’-CGTGGAATGCGTTCAGTTGG-3’,

LETM1_reverse (rev) 5’-GAAGCTGATGGAGGTAATTGAG-3’; Nanos-1_fw 5’-ACTTGTCCATTATGCGGTGCT-3’, Nanos-1_rev 5’-GGTTCCAACAAACCAGCTTCA-3’; Bcl2_fw 5’-TCTTCATGATGGTTGGTCTGGA-3’, Bcl2_rev 5’CCGACAAGAGCAGCTAAACC-3’; FGFR1_fw 5’-CACAGAAGGAGATGTGTCTGAA-3’ FGFR1_rev 5’-TTCCCGTAAGGAGCATATTCCA-3’.

### Statistical Analysis

The data were analyzed using the software Prism 10, Version 10.1.2 (GraphPad Software, LLC, San Diego, CA, USA). Quantitative data are expressed as mean ± SEM and comparative analysis among was conducted using one-way and two-way ANOVA followed par multiple comparison tests (> 2 groups) or Mann-Whitney U test (= 2 groups). Statistical significance was accepted when *p* < 0.05.

## Results

### Fractionation of mouse serum reveals PAFAH as an active anti-schistosomal multi-stage enzyme

Humans are known to be the primary definitive host for *S. mansoni*, while the mouse is the most widely used experimental laboratory host. Recently, we observed that when human serum (HSe) was replaced by mouse serum (MSe) in our cell-free *ex vivo*-culture system for molecule and drug testing (27, 28), newly transformed schistosomulae (NTS) as well as developing advanced larval stages died within 72 hours (10), suggesting that MSe has soluble serum factor(s), which are detrimental for survival and development of mammalian host dwelling stages of *S. mansoni* (27, 28). To identify the schistosomicidal serum factor(s), we performed a large-scale fractionation of MSe using a Superdex 200 size exclusion column, pooled 35 fractions into 5 subfractions (F1-5), and tested these in the NTS bioassay (Fig. 1A and Supplementary Fig. S1A). Our *ex vivo* study revealed two fractions (F1 and F2) with NTS killing effect, whereby only fraction F2 very efficiently killed NTS comparable to the schistosomicidal effect of 20% MSe (Fig. 1B). Furthermore, heating (99°C), but not Pronase E treatment of MSe and fraction F2, reduced and reliably abolished NTS killing capacity (Supplementary Fig. S1B), indicating that the NTS killing effects could be due to a protein, including lipoproteins and phospholipoproteins. Next, to identify the protein(s) responsible for the killing activity, we analyzed the different killing and non-killing fractions using comparative mass spectrometry. Afterward, the subtraction of proteins found in the inactive fractions and heated fractions from the set of proteins identified in the active fraction F2 allowed the restriction of the effective proteins in an initial step to 69 potential candidates for the active killing component in MSe (Supplementary Fig. S1C).

**Figure 1.**
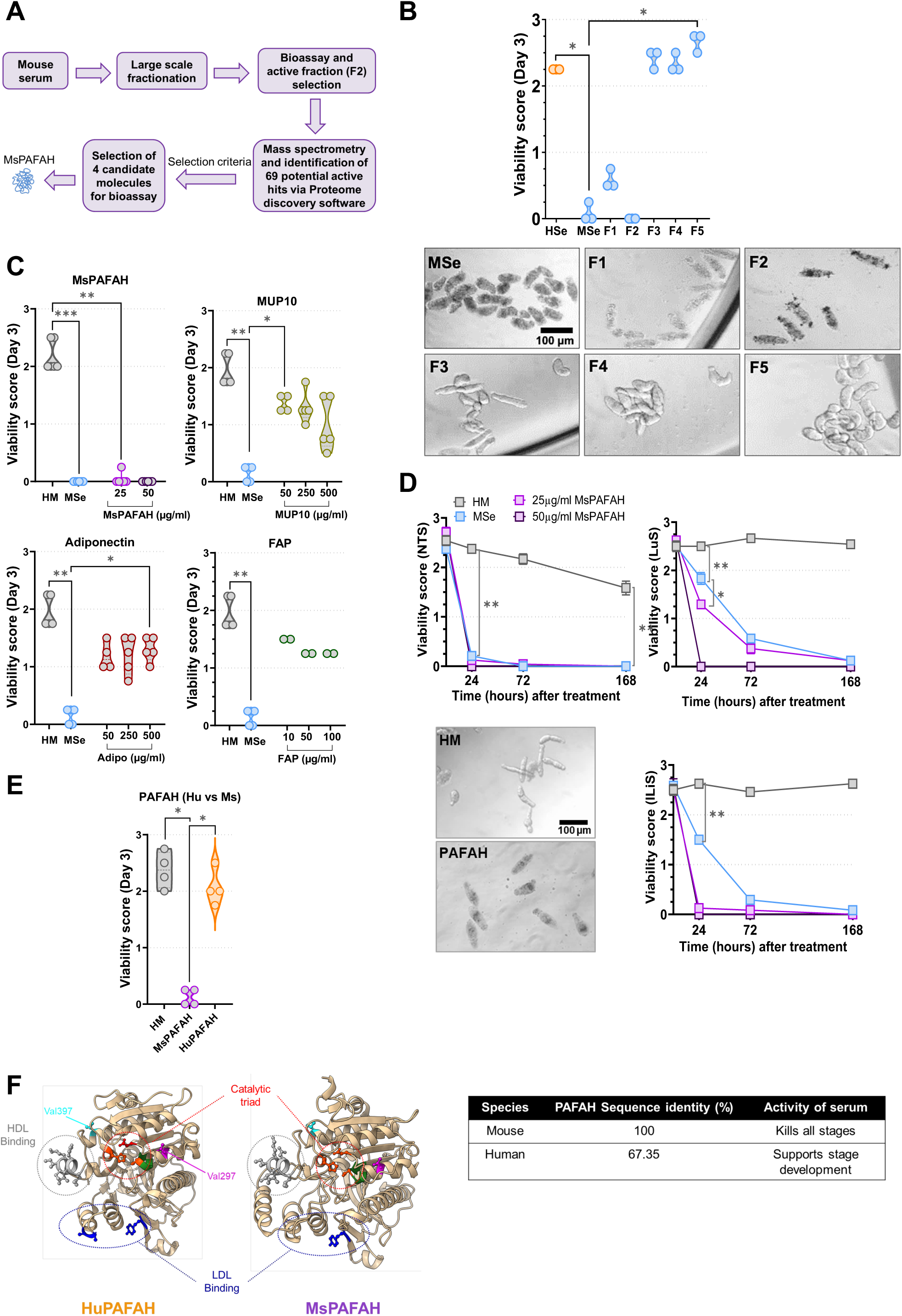
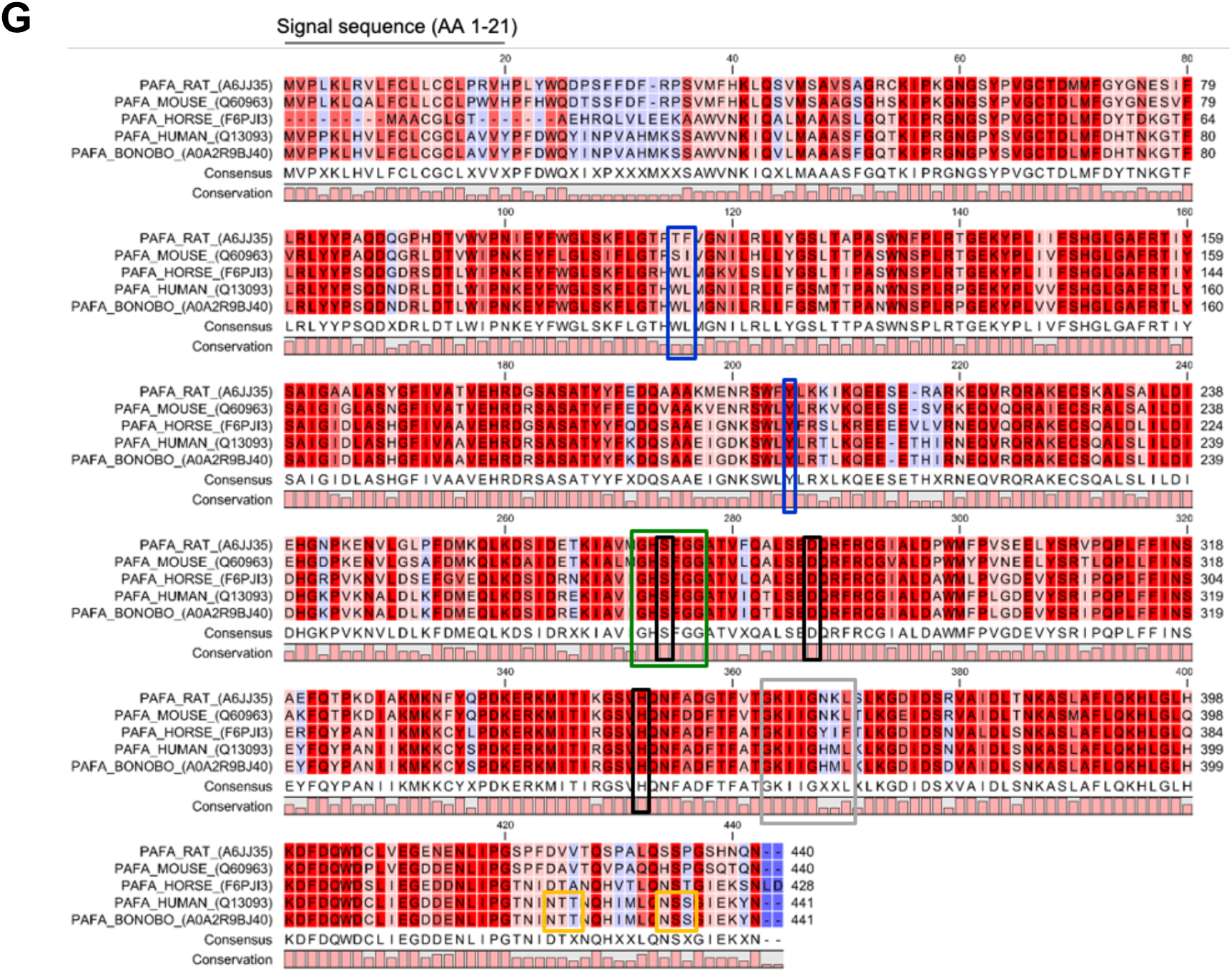
Large-scale fractionation of mouse serum reveals MsPAFAH as an active anti-schistosomal multi-stage enzyme. (A) Schematic procedure of large-scale fractionation of mouse serum and identification of candidate molecules and mouse platelet-activating factor-acetylhydrolase (MsPAFAH). (B) Effect of human (HSe) and mouse (MSe) sera and fractions at 20% on the viability of newly transformed schistosomulae (NTS) at day 3 (left panel). A representative picture is shown on the right panel. (C) Concentration-dependent effect of mouse serum selected recombinant expressed candidate molecules, MsPAFAH, major urinary protein 10 (MUP10), fibroblast activation protein (FAP) and Adiponectin as compared to mouse serum (MSe) and HM control on NTS at day 3. (D) Time-dependent effect of MsPAFAH on different *S. mansoni* developmental larval stages (skin (NTS), lung (LuS), and late liver juvenile (lLiS)). (E) Comparison of the effect of recombinant mouse (MsPAFAH) and human PAFAH (HuPAFAH) on NTS at day 3. (F) Structural comparison and homology modeling of HuPAFAH and MsPAFAH. Both structures have a classic α/β serine hydrolase fold and contain a catalytic triad (Ser273, His351, and Asp296 in HuPAFAH; Ser272, His350, and Asp295 in MsPAFAH (in red) and HDL/LDL (Low/high-density lipoproteins) binding sites (blue) (upper panel) with sequence identity and killing properties (lower panel). (G) Alignment of selected PAFAH amino acid sequences (HUGO gene name: PLA2G7). Sequences were retrieved from UniProt (rat: A6JJ35; mouse: Q60963; human: Q13093; bonobo: A0A2R9BJ40; horse: F6PJI3) and aligned using Clustal Omega default settings. Data information and statistics: Results are representative of at least three independent experiments and are expressed as means ± SEM. Asterisks show significant statistical differences analyzed using Kruskal–Wallis one-way ANOVA followed by a Dunn’s multiple comparison test (B, C, E) and two-way ANOVA followed by Tukey’s multiple comparison test (D). *P < 0.05; **P < 0.01; ***P < 0.001.

From these 69 potential candidates, four (PAFAH, major urinary protein 10 (MUP10), fibroblast activation protein (FAP), and adiponectin) were chosen for recombinant expression based on different criteria as detailed in the methods. All four purified candidate proteins were tested for their effect on the viability of NTS as compared to MSe and HSe (Fig. 1C). Interestingly, mouse PAFAH (MsPAFAH) had the strongest viability-reducing effect on NTS after 72 h with a significant impact visible at a concentration of 15 µg/mL with a lethal effect at 25 µg/mL (Fig. 1C, Supplementary Fig. S1D). The enzymatic activity of the recombinant MsPAFAH was confirmed using a commercial assay and, although used at higher concentration, was less active when compared to PAFAH activity of MSe correlating with the slightly lower killing efficiency *in vitro* (Supplementary Fig. S1E) as compared to MSe. We also observed killing properties for the other proteins, however, at comparably higher concentrations of 500 µg/ml (Fig. 1C) with no effect at lower concentrations (less than 10 µg/ml) (Supplementary Fig. S1F). Thus, we moved on with investigating in more detail MsPAFAH, which showed the closest schistosomicidal profile with mouse serum. We next investigated the killing impact of MsPAFAH on more advanced *in vitro*-generated *S. mansoni* larval stages (lung stage (LuS), late liver juvenile (lLiS)), critical for host reinfection. As for MSe, almost all mammalian host-dwelling stages of *S. mansoni* were killed within 72 h (Fig. 1D). Of note, LuS was less affected by MSe and MsPAFAH at 72 h as compared with NTS and lLiS, the closest developmental stage to the adult worm in human schistosomiasis (Fig. 1D). Interestingly, the killing efficiency of MsPAFAH of NTS was strongly impaired in presence of the selective secretory phospholipase A_2_ inhibitor Vareplasdib (VAR) (33), which, however, was less pronounced when preincubating VAR with MSe (Supplementary Fig. S1G), indicative of potential further killing component(s) in MSe.

Since humans are the major definitive host of *S. mansoni*, we next investigated the impact of recombinant human PAFAH (HuPAFAH) on the viability of the mature worm. Interestingly, we observed no killing effect by HuPAFAH as compared to MsPAFAH (25 µg/mL each) (Fig. 1E), reflecting our previous findings that HSe did not only fail to kill NTS but also promoted their development *in vitro* to juvenile worms (10). Importantly, sequence alignment and homology-modeling of HuPAFAH and MsPAFAH structures (46, 47) (48) revealed that while the Ser/Asp/His catalytic triad of serine hydrolases as well as other residues (Val297 and Val397) important for activity are perfectly conserved, there are major differences in the LDL binding region (Fig. 1F, Fig. 1G), relevant for the enzymatic activity. In addition, their amino acid sequences showed only 67.35% overall identity, including the 21 amino acid signal sequences (Fig. 1F, Fig. 1G).

Collectively, our data so far indicate that MsPAFAH represents one of the strongest schistosomicidal components present in MSe and is responsible for the efficient killing of different stages of *S. mansoni ex vivo*.

### PAFAH is upregulated during *S. mansoni* infection in mouse serum and reduces the reproductivity of adult worms

To investigate whether PAFAH is regulated *in vivo* during schistosome infection as an indication of its potential role in host defense, we infected mice for 7 weeks with *S. mansoni* and evaluated serum levels of PAFAH in infected and naïve control animals. As depicted in Fig. 2A, the infection markedly increased MsPAFAH levels in the serum. Having shown that MsPAFAH profoundly influences the viability of *ex vivo*-generated *S. mansoni* developmental stages and is upregulated during infection, we examined the impacts of MSe and MsPAFAH on adult worms isolated from infected mice. For this, mature *S. mansoni* worms of both sexes were flushed from infected mice and cultured *ex vivo* in couples or individually (Fig. 2B-C). Interestingly, MSe and MsPAFAH similarly reduced the viability of worm couples in a time- and concentration-dependent manner, with a significant effect of MsPAFAH at 15 µg/mL within 48-72 h (Fig. 2B). Importantly, although both sexes of *S. mansoni* were heavily affected after 72 h of treatment with MSe and MsPAFAH, we observed a sex-biased viability effect with males being more susceptible to the treatment (Fig. 2C). Similar sex-differences were previously reported for schistomicidal small molecules, such as PZQ (49, 50).

**Figure 2.**
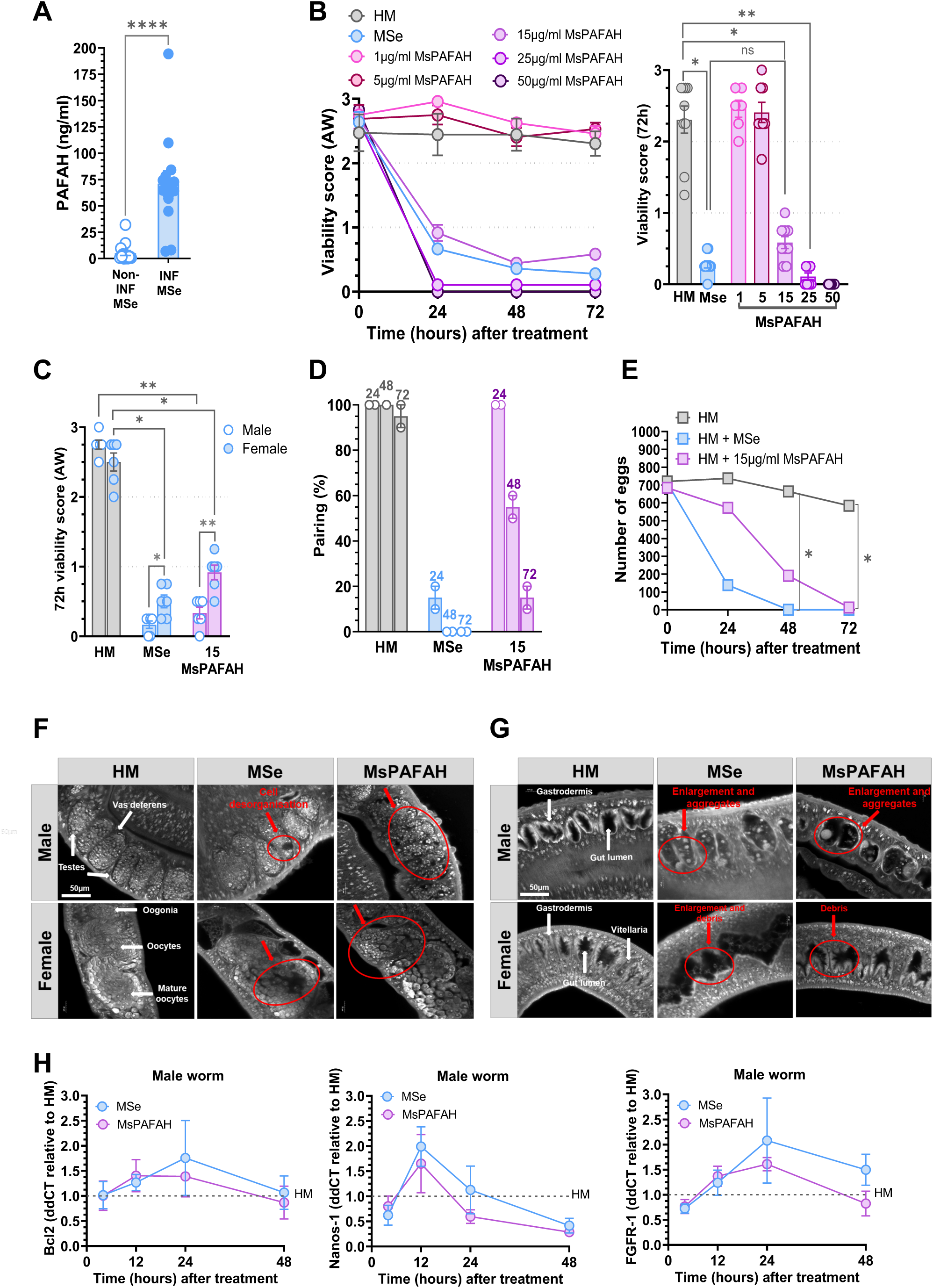
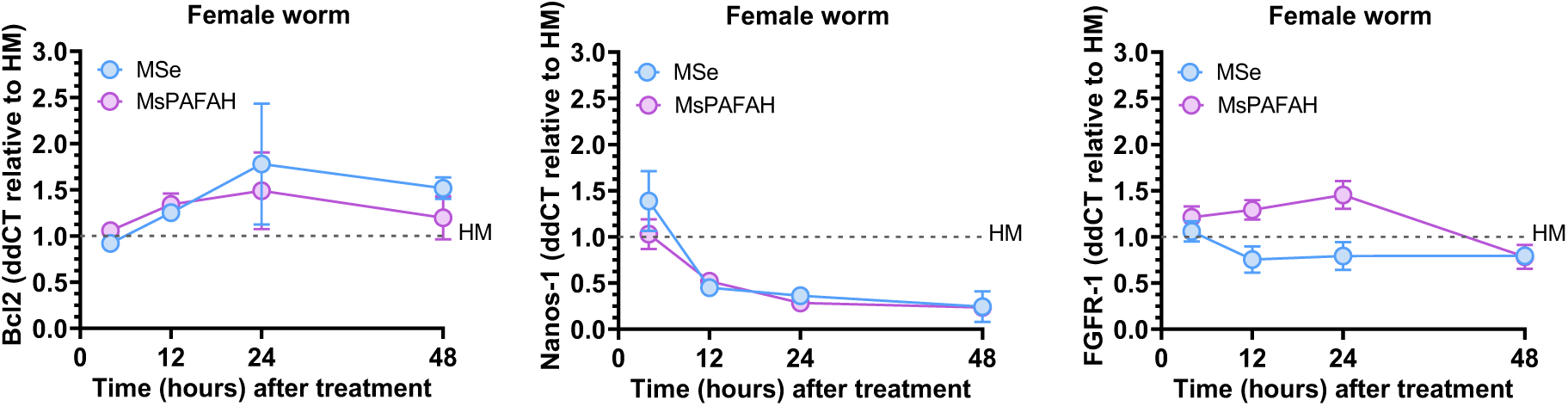
MsPAFAH is upregulated during *S. mansoni* infection in serum and reduces the reproductivity of adult worms. (A) Serum levels of MsPAFAH in schistosome-infected and non-infected mice. (B) Concentration- and time-dependent effects of MsPAFAH on ex-vivo adult worms recovered from infected mice. (C) Effects of MsPAFAH and MSe on male compared to female adult worms at day 3. (D) Impact of MsPAFAH and MSe on male and female adult worm pairing. (E) Number of eggs released by female adult worms over time following treatment with MsPAFAH and MSe. (F) CLSM-analysis of the effect of MsPAFAH and MSe treatment on gonadal tissues of male and female adult worms at day 3. White arrows show structure localization. Red arrows show affected areas and tissue disorganization by MsPAFAH and MSe. (G) CLSM-analysis of the effect of MsPAFAH and MSe treatment on intestinal tissues of male and female adult worms at day 3. White arrows show structure localization. Red arrows show affected areas and aggregate and debris formation by MsPAFAH and MSe. (H) qPCR analysis of MsPAFAH- and MSe-driven regulation of the expression of anti-apoptotic and gonadal stem cell proliferation- and differentiation-related genes in male and female adult worms. Data information and statistics: Results are representative of at least three independent experiments (5-10 worm pairs per condition) and are expressed as means ± SEM. Asterisks show significant statistical differences analyzed using Kruskal–Wallis one-way ANOVA followed by Dunn’s multiple comparison test (A, B, C) and two-way ANOVA followed by Tukey’s multiple comparison test (E). *P < 0.05; **P < 0.01; ****P < 0.0001.

Human schistosomiasis mainly results from the immunopathology induced by host responses to deposited eggs in tissues, such as in the liver (51, 52). Unlike other trematodes, successful egg production depends on worm pairing stability (51, 52). We thus conducted a pairing stability assay in the presence of MSe and MsPAFAH (Fig. 2D). We observed an 80% reduction of pairing stability at 24 h post-treatment (p. t.)with MSe. As expected, MsPAFAH also affected the pairing stability of *S. mansoni*; however, the effect was a bit delayed (Fig. 2D). Accordingly, we noted a significant reduction in the number of eggs deposited in culture at 48 h post-treatment for both MSe and MsPAFAH, with a complete arrest in egg production at 72 h compared to the non-treated control in hybridoma medium (HM) (Fig. 2E). To investigate whether these observations were related to morphological defects in the parasite’s reproductive system, MSe- and MsPAFAH-treated worms were fixed for subsequent confocal laser scanning microscopy (CLSM) analysis. We detected morphological abnormalities at the organ level (Fig. 2F-G). In untreated control males, testes were composed of several testicular lobes containing numerous spermatogonia and spermatocytes in different stages of maturation (Fig. 2F, upper panel left). Physiologically, spermatogenesis is located at the dorsal part of the lobes with large round spermatogonia and expires at the ventral part, indicated by smaller elongated mature sperms (spermatozoa) (37, 38). After MSe and MsPAFAH treatment, these lobes appeared disorganized and showed porous areas, creating a Swiss*-*cheese-like tissue pattern (Fig. 2F, upper panel, middle and right). Nevertheless, the seminal vesicle still contained some spermatocytes of unknown quality. In control females, the small immature oogonia were detected within the anterior part of the ovary, and bigger mature oocytes were found within the larger posterior part (37, 38). We observed that MSe and MsPAFAH also disrupted the structure of the ovaries (Fig. 2F, lower panel, middle and right). They appeared disorganized and shrunken, and oogonia of smaller size and mature oocytes were poorly separated from each other. Unlike control ovaries, hole-like areas were found within the ovary, indicating a lack of cell-based staining. In contrast, in the absence of MSe and MsPAFAH treatment, the vitellarium of control females was found to be tightly packed with cells, which interlocked with the cell rows of the opposite side in a zipper-like arrangement as previously reported (37, 38).

We further evaluated the internal structure damage of the digestive organs of treated worms (Fig. 2G). Whereas the gastrodermis of control males appeared as a thick, continuous layer of syncytial cells (53, 54), MSe and MsPAFAH-treated worms showed an enlarged gut lumen and degradation of the gastrodermis, which led to the accumulation of degraded tissue and particle aggregates of remarkable size (Fig. 2G, upper panel, middle and right). Especially, the gastrodermis appeared to be detached from the adjacent tissue layer and completely collapsed in some compartments. Although females appeared less affected by the internal destruction of the digestive organs, reflecting the viability assessment, enlargement and debris were noted following MSe and MsPAFAH treatment (Fig. 2G, lower panel, middle and right).

In summary, treatment with MSe and MsPAFAH led to a dramatic reduction of schistosomal eggs. This might be related to the structural disruption following apoptotic signaling and disintegration of stem and reproductive cells in the ovary and testes, as reported elsewhere (55, 56). Therefore, we investigated the regulation of genes related to apoptosis (*bcl2*) and gonadal stem cell proliferation, stability, and differentiation (*nanos1, fgfr1*) important in female reproduction (55, 56). In line with the degraded gonadal cells noted by CLSM, we observed a downregulation of the expression of *nanos1* and, to a lesser extent, *fgfr1* in both male and female adult worms in the presence of MSe and MsPAFAH (Fig. 2H). In contrast, there was an increase in the expression of anti-apoptotic *bcl2* after 12-24 h following MSe and MsPAFAH treatment, which remained after 24 h and 48 h only in female worms, possibly as a sign of better survival and active apoptosis regulating mechanisms as compared to males (Fig. 2H). Of note, except for *fgfr1* regulation in female worms, both MSe and MsPAHAH had comparable effects overall.

Taken together, these data suggest that, in mice, PAFAH might be a potent host defense enzyme regulated during schistosome infection with the capacity to severely reduce the parasite viability and reproductive activity.

### MsPAFAH manipulates the schistosome phosphatidylethanolamine (PE) and phosphatidylcholine (PC) composition and distribution

Having characterized the internal structural degradation induced by PAFAH present in MSe, we next investigated whether the observed phenotypes could originate from an effect on schistosomal surface membranes and potential apoptotic signaling as reflected by massive granulation in parasite tissues in the presence of MsPAFAH and MSe. Thus, we conducted scanning electron microscopy (SEM) imaging of formalin-fixed adult worms following treatment with MSe and MsPAFAH. Compared to the untreated HM control, we observed, in the treated groups, contracted worms at lower magnification (Fig. 3A, upper panel, middle and right) and, at higher magnification, damaged and disorganized tegumental structures at 72 h p. t. (Fig. 3A, lower panel, red circles).

**Figure 3.**
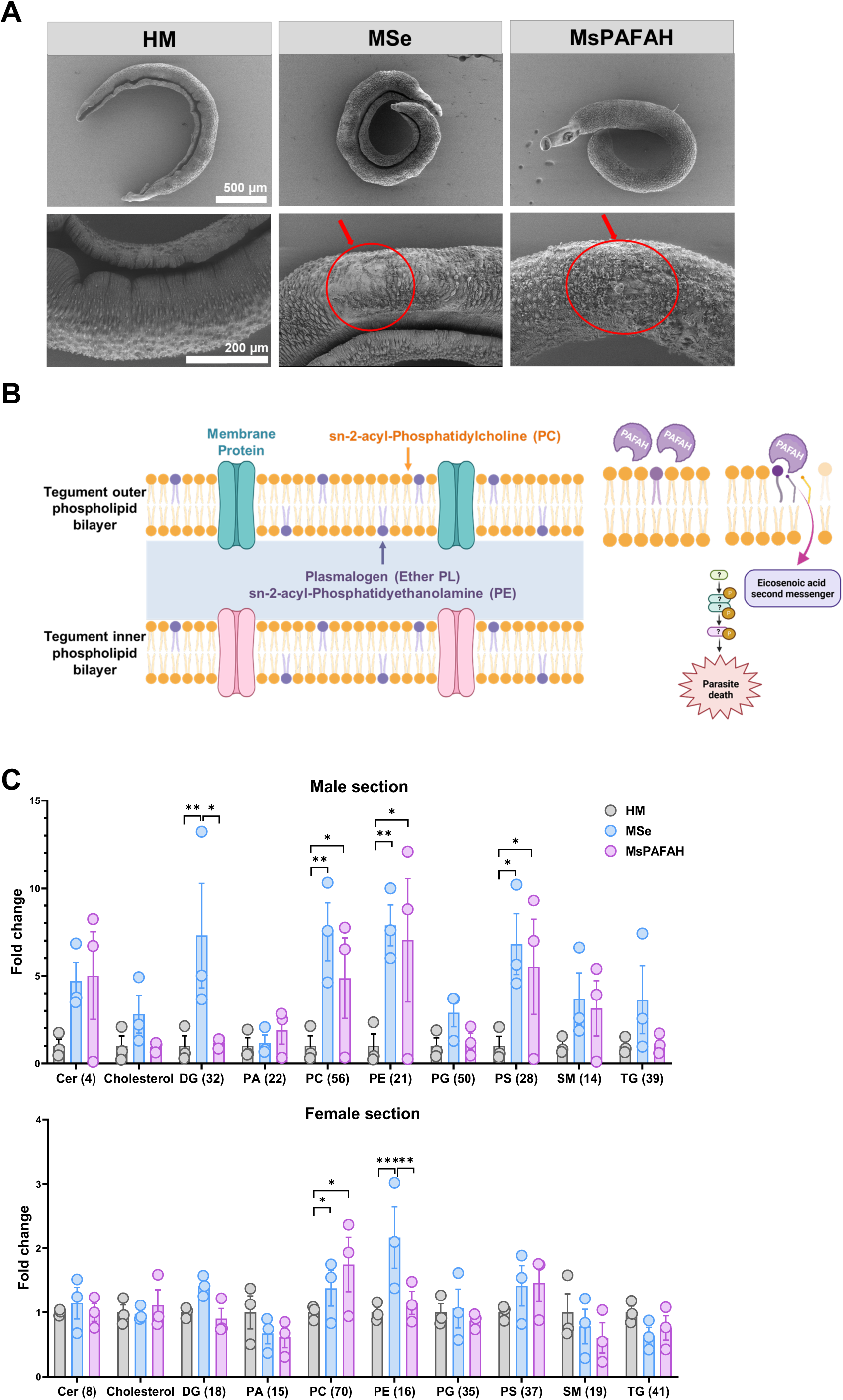
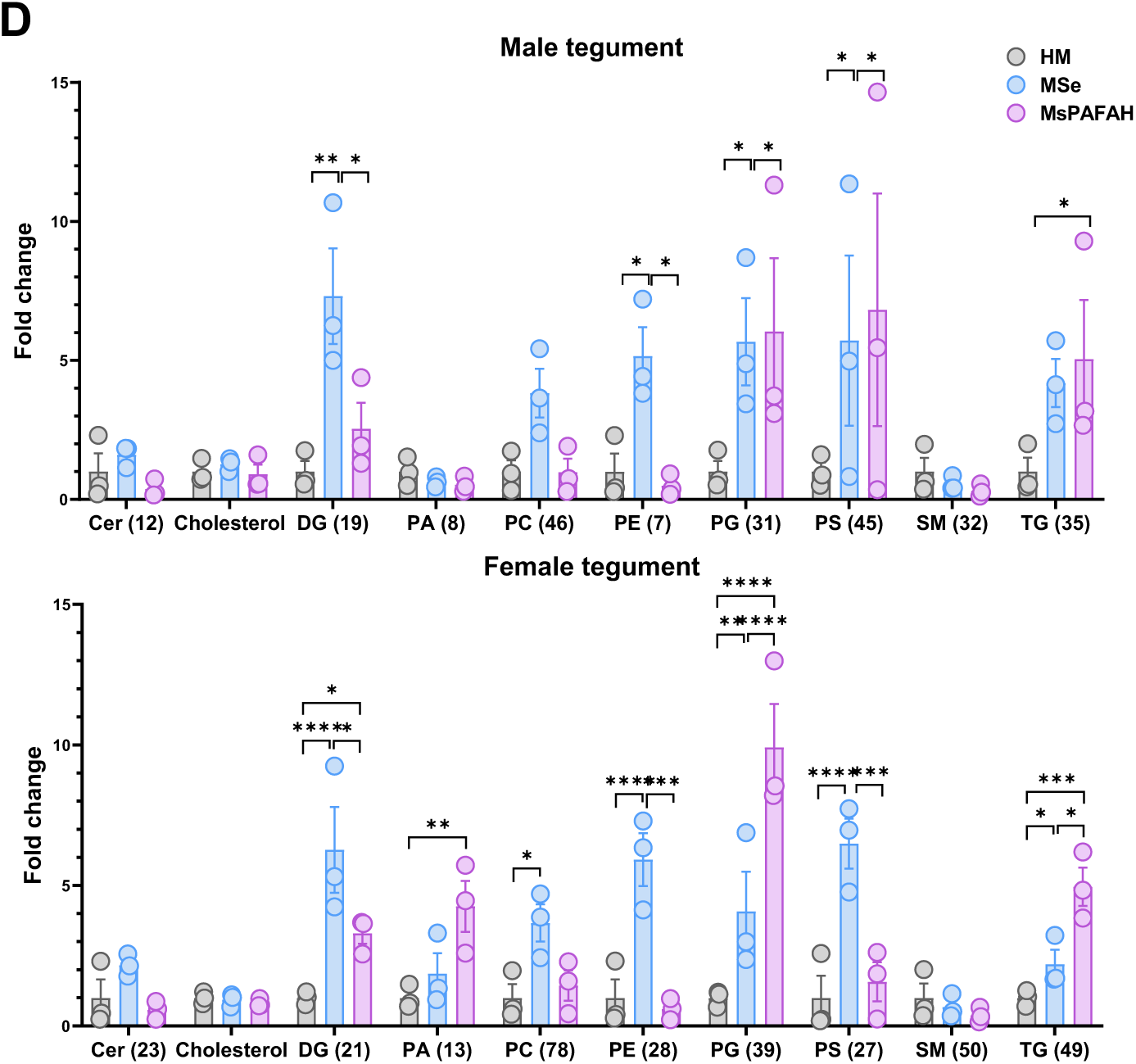
MsPAFAH remodels the schistosome phospholipids PE and PC composition and distribution. (A) SEM analysis of the effect of MsPAFAH and MSe treatment on ex-vivo adult worm tegument at day 3. Red arrows show disorganized tegument tissue layers following MsPAFAH and MSe treatment. (B) Schematic representation of schistosome tegumental phospholipid layers (left panel) and investigation hypothesis of the effect of MsPAFAH and potential downstream signaling. C) MALDI-MSI analysis of fold change in the composition of male (upper panel) and female (lower panel) adult worm phospholipids and lipids following MsPAFAH and MSe treatment as compared to HM control. Numbers in brackets are the number of lipids detected and analyzed. Cer: Ceramides; DG: Diacylglycerols; PA: Phosphatidic acids; PE: Phosphatidylethanolamines PC: Phosphatidylcholines; PG: Phosphatidylglycerols; PS: Phosphatidylserines; SM: Sphingomyelins; TG: Triglycerides. (D) MALDI-MSI analysis of MsPAFAH- and MSe-induced fold change of the major male (upper panel) and female (lower panel) schistosome tegument phospholipid classes. Numbers in brackets are the number of lipids analyzed. Data information and statistics: Results are representative of at least two to three independent experiments (5 worms) and are expressed as means ± SEM. Asterisks show significant statistical differences analyzed using Kruskal–Wallis two-way ANOVA followed by Tukey’s multiple comparison test. *P < 0.05; **P < 0.01; ***P < 0.001.

The tegument of schistosomes is essential for the survival of the parasite (50). As presented in Fig. 3B, the tegument of *S. mansoni* is composed of a syncytial structure consisting of two (outer and inner) closely apposed phospholipid (PL) bilayers that form the interactive surface with the host. PAFAH hydrolyses PL with short-chain acyl groups (n<8) at sn-2 position (sn-2-acyl-PLs e.g. platelet-activating factor (PAF)) as well as oxidized PLs with longer acyl chains having aldehyde, carboxylic acid, hyperperoxid or epoxy groups, thereby generating lyso-PAF, lyso-PLs (57), free fatty acids and lipid mediators (58). In addition, PAFAH has a transacetylase activity contribtuting to PAF diversity (58). Therefore, we hypothesized that MsPAFAH binds to, hydrolyzes, and disrupts worm tegument sn-2-acyl-PLs, leading to worm death, possibly involving oxidized PLs in schistosomes, whose presence is controversial(59). We thus first analyzed the lipid profile of male and female worm sections (Fig. 3C) and teguments (Fig. 3D) using untargeted metabolipidomics and imaging. The matrix-assisted laser desorption/ionization mass spectrometry imaging (MALDI-MSI) surprisingly revealed an upregulation of PL levels in response to both MSe and MsPAFAH after 72 h of treatment, both in the body (Fig. 3C) and in the tegument (Fig. 3D) of the worms, with a significant change of PE and PC, the major PLs in schistosomes (50) and less abundant lipids such as the triglycerides (TG).

These data demonstrate a remodeling of the distribution and composition of PLs, mainly of the PE and PC species, upon MsPAFAH and MSe treatment.

### MsPAFAH, but not HuPAFAH, depletes specific unsaturated PC and PE species of schistosomes while increasing overall diacyl- and lyso-phospholipid levels

To gain more insights into how MSe and MsPAFAH remodel adult worm PC and PE species, we performed quantitative metabololipidomics using a more sensitive ultra-performance liquid chromatography tandem mass spectrometry (UPLC-MS/MS) system (59, 60). Profiling of PC and PE species revealed that male worms have a higher level of total diacyl-PC and diacyl-PE than female worms (Supplementary Fig. S2A), which was expected since male worms are larger. In addition, the fatty acid compositions in diacyl-PC and diacyl-PE significantly differ between male and female worms (Supplementary Fig. S2B). For example, the dominant PC species PC(palmitate (16:0)_eicosenoate (20:1)) was considerably higher in male than in female worms (42.3% vs. 26.4%). In contrast, higher levels of PE(stearoyl (18:0)_adrenate (22:4)) were found in females (42.4% vs 36.3%) (Supplementary Fig. S2C). These data confirmed baseline sex differences in membrane PL profile in *S. mansoni* adult worms reported previously (50).

We next sought to identify the effect of MSe and MsPAFAH on specific membrane PLs. Interestingly, male worms responded stronger to MSe- and MsPAFAH treatment than female worms (Fig. 4A-B, left panels). MSe significantly elevated total diacyl-PC levels in male worms, whereas MsPAFAH, rather, markedly increased total diacyl-PE levels (Fig. 4A-B, left panels). By trend, MSe and MsPAFAH also slightly elevated the total diacyl-PC and diacyl-PE content in female worms. Principle component analysis of the diacyl-PC or diacyl-PE profiles revealed that the treatment with MSe and MsPAFAH did not shift the diacyl-PC or diacyl-PE compositions in similar directions (Fig. 4A-B, lower panels). Therefore, we further explored potential changes in individual diacyl-PC or diacyl-PE species or subgroups modulated by both MSe and MsPAFAH. MSe or MsPAFAH treatment substantially altered the availability of various individual diacyl-PC (Fig. 4C) and diacyl-PE species in male and female worms (Fig. 4D) overall. We displayed the diacyl-PC and diacyl-PE species in volcano plots to identify PLs, whose levels are changed by both MSe and MsPAFAH in the same direction and thus point to substrates or lipids specifically regulated by MsPAFAH in MSe. Further detailed analyses are shown in Supplementary Fig. S3 and Supplementary Fig. S4. The differentially regulated species of interest were selected using the following criteria: 1) the relative proportions (% of total diacyl-PC or diacyl-PE) > 0.5%, 2) fold changes against HM control ≤ 0.9 or ≥ 1.1, and 3) p values (MSe or MsPAFAH against HM control) < 0.1. We identified a small number of shared diacyl-PE or diacyl-PC species, whose levels were comparably altered by both MSe or MsPAFAH treatments in male and female worms (Fig. 4C-D). Of interest, mapping the differentially regulated PL species highlighted PC(16:0_20:1) and PE(18:0_22:4) (Fig. 4C-D), the most abundant diacyl-PC or diacyl-PE species, to be significantly decreased by both MSe and MsPAFAH treatment regardless of the gender of the worms (Fig. 4E-F). On the other hand, the proportions of PC(18:0_18:2) (Fig. 4E) and PE(18:0_18:2) (Fig. 4F) were elevated by both treatments, as also confirmed by MALDI-MS imaging (Fig. 4E-F, right panels). Of note, PC(18:0_18:2) and PE(18:0_18:2) comprise only 1-2% of total diacyl-PC or diacyl-PE, which was much lower than PC(16:0_20:1) and PE(18:0_22:4) (30-50%) (Fig. 4E-F). Nonetheless, this effect on these PC and PE species is only observed with MsPAFAH, but not in the presence of HuPAFAH (Fig. 4G). Interestingly, when looking at hydrolysis products of PC(16:0_20:1) and PE(18:0_22:4) using MALDI-MS imaging, we noticed an increase of lyso-PC(20:1) and lyso-PE(18:0) (Fig. 4H), in line with the decline of the respective MsPAFAH-hydrolysed PC and PE.

**Figure 4.**
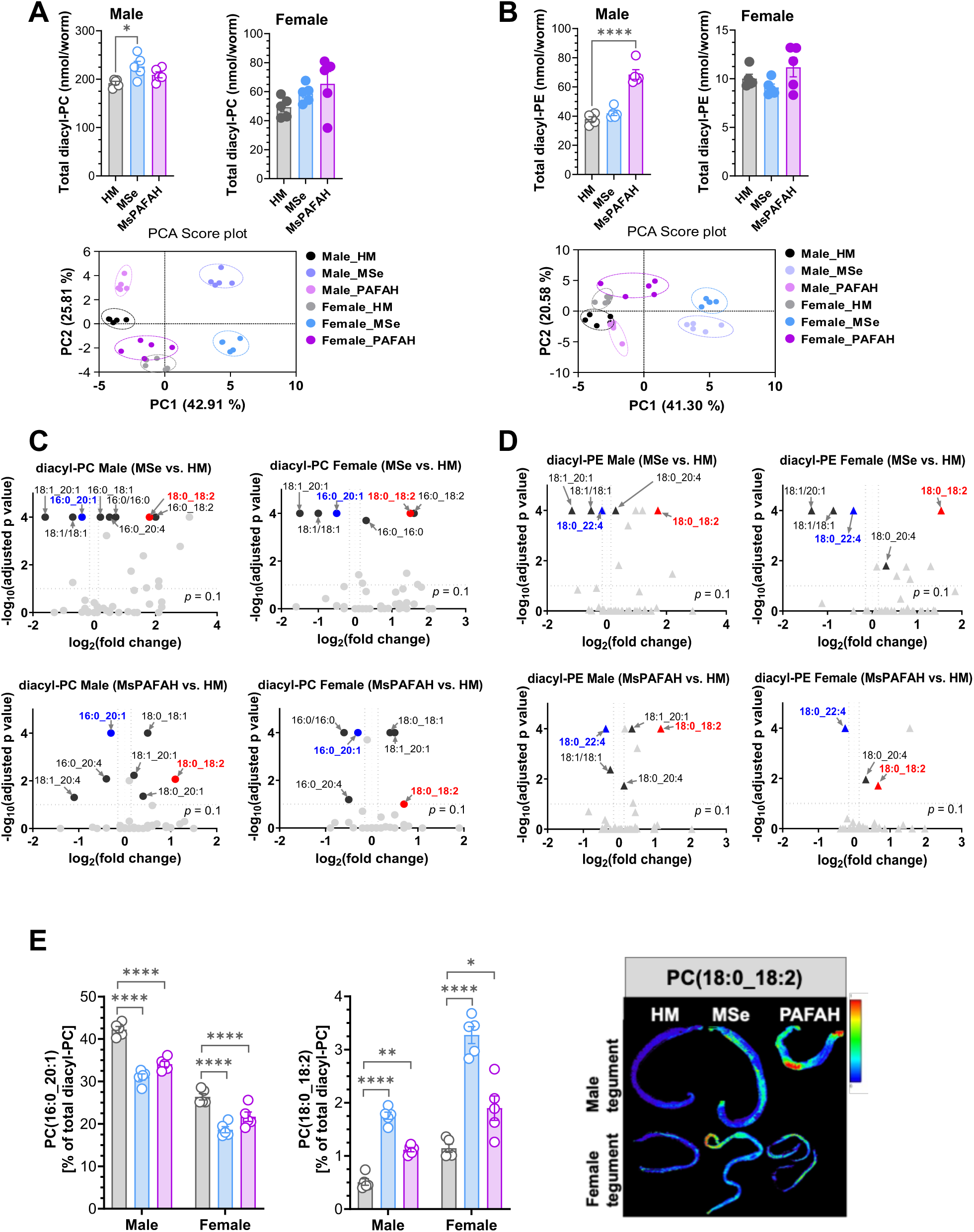
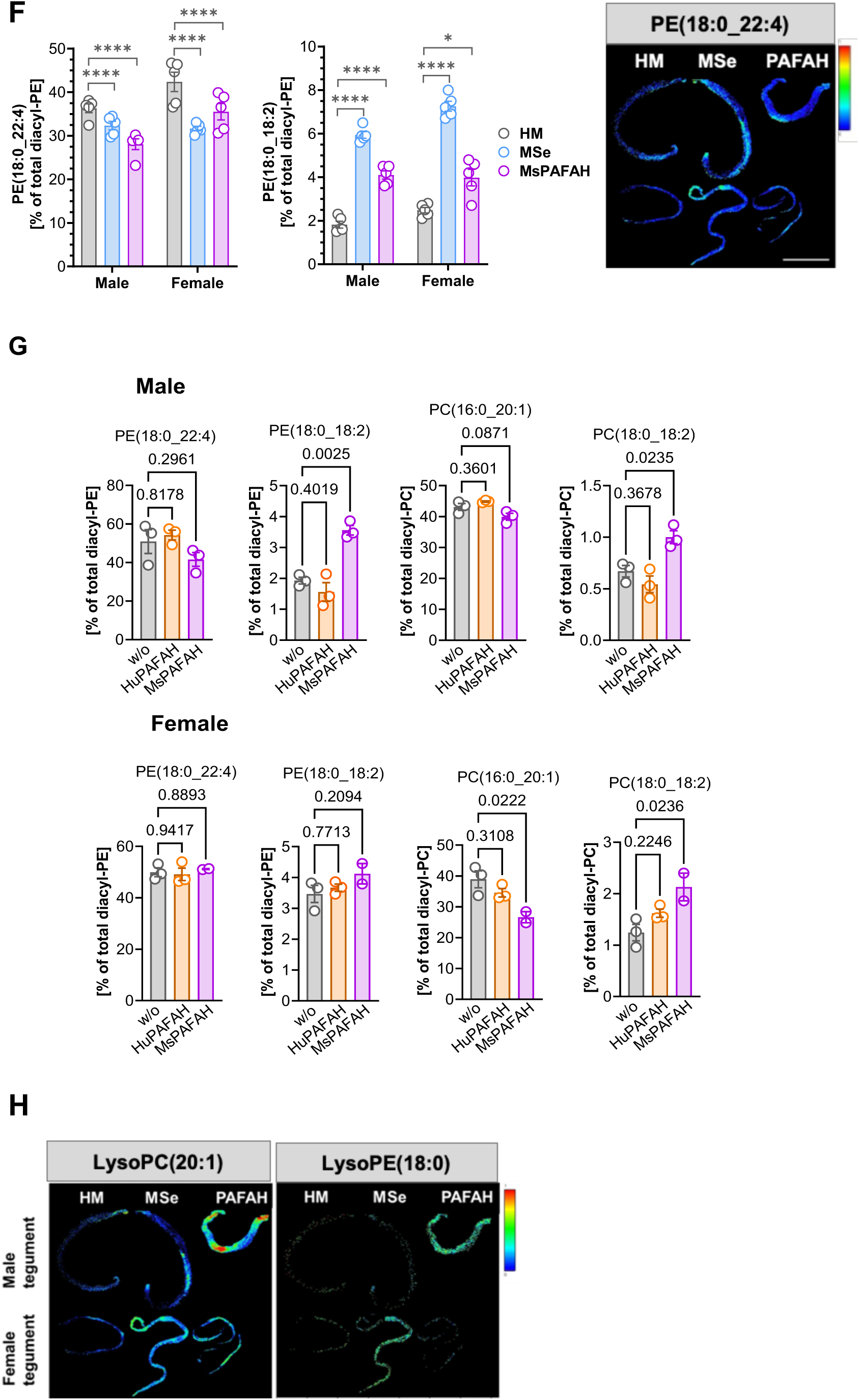
MsPAFAH, but not HuPAFAH, depletes specific unsaturated PC and PE species of schistosomes while increasing overall diacyl- and lyso-phospholipid levels. (A) UPLC-MS/MS analysis of extracted diacyl-phosphatidylcholine (PC) from male and female adult worms (upper panel) and PCA score plots with clustering (lower panel) per sex following MsPAFAH and MSe treatment. Graphs show the total diacyl-PC amount (pmol/worm, upper panel); PCA plots were generated using the relative intensities of individual diacyl-PC species (% of total diacyl-PC). (B) UPLC-MS/MS analysis of extracted diacyl-phosphatidylethanolamine (PE) from male and female adult worms (upper panel) and PCA score plots with clustering per sex (lower panel) following MsPAFAH and MSe treatment. Graphs show the total PE amount (pmol/worm, upper panel); PCA plots were generated using the relative intensities of individual diacyl-PC species (% of total diacyl-PC). (C) Volcano plots showing the differentially regulated diacyl-PC species (relative intensities, % of total diacyl-PC) upon MsPAFAH and MSe treatment in male and female worms. (D) Volcano plots showing the differentially regulated diacyl-PE species (relative intensities, % of total diacyl-PE) upon MsPAFAH and MSe treatment in male and female worms. (E) Proportion of the downregulated PC(16:0_20:1) and upregulated PC(18:0_18:2) species following MsPAFAH and MSe treatment. (F) Proportion of the downregulated PE(18:0_22:4) and upregulated PE(18:0_18:2) species following MsPAFAH and MSe treatment. The right panels show MALDI-MS imaging of the PC (18:0_18:2) and PE(18:0_22:4) species. (G) Proportion of the downregulated PC(16:0_20:1) and PE(18:0_22:4), and upregulated PC and PE(18:0_18:2) species following MsPAFAH as compared to HuPAFAH treatment and control. (H) MALDI-MS imaging of the MsPAFAH-hydrolyzed products of PC(16:0_20:1) and PE(18:0_22:4). Data information and statistics: Results are representative of at least three independent experiments (5 worms in each group) and are expressed as means ± SEM. For panels A, B, C, D, E, F, and G, statistical p values were calculated by ordinary one-way (A, B, G) or two-way (C, D, E, F) ANOVA + Dunnett’s post hoc test against HM-treated worms, *P < 0.05; **P < 0.01; ***P < 0.001; ****P < 0.0001. For panels C and D, fold changes were calculated from the relative intensities (% of total diacyl-PC or diacyl-PE) of the species from MsPAFAH- or MSe-treated against HM-treated worms. The differentially regulated species were marked when the following criteria were met: 1) the proportion of the species (% of total PC or PE) > 0.5%, 2) fold changes against HM control ≤ 0.9 or ≥ 1.1, 3) P values < 0.1 adjusted.

In summary, MSe and MsPAFAH strongly impact the PL composition of *S. mansoni* by reducing abundant diacyl-PC and diacyl-PE species and increasing the hydrolysis products. Potentially, in an adaptive response to this dysregulation, the total amount of schistosomal diacyl-PL, in particular, 18:2-containing species, is substantially upregulated.

### PAFAH remodels schistosomal ether PE abundance and composition in a sex-dependent manner

The substrate specificity of plasma PAFAH is not limited to PAF, diacyl-PC, and diacyl-PE but also includes ether- and oxidized PLs (57, 61). In fact, plasmanyl and plasmenyl PE (with an ether-bound fatty alcohol at the sn-1 position and an acyl group at the sn-2 position) constitute a substantial fraction of the *S. mansoni* tegument membrane PLs (26, 50). As with total diacyl-PE and diacyl-PC, overall baseline levels of ether PE were higher in males as compared to female worms, similarly for plasmenyl PE and plasmanyl PE (Fig. 5A-B). However, although no differences were seen for total ether PE and plasmenyl PE following exposure to MsPAFAH and MSe, we observed an increase of plasmanyl PE levels in adult male worms in contrast to females (Fig. 5A-B). Furthermore, MSe and MsPAFAH inversely altered the ratio of diacyl-PC, diacyl-PE, and ether PE (Fig. 5C) in male and female worms, thereby revealing gender differences due to treatment which are not present at baseline in untreated worms (Fig. 5C). We further analyzed whether MSe and MsPAFAH affect the proportions of individual ether PE species in male and female worms. Overall, MSe was more effective than MsPAFAH in remodeling the ether PE composition. Of note, shared effects of MSe and MsPAFAH between males and females are mainly the decrease in the proportion of PE(P-16:0/18:1) and the increase in the proportion of PE(P-16:0/20:4) (Fig. 5D upper and lower panels). The effects were more pronounced in males than in females, as shown by the example of PE(P-16:0/18:1) (Fig. 5E, left panel) and PE(P-16:0/20:4) (Fig. 5E, right panel). Importantly, the stronger effect of MSe, as compared to MsPAFAH, in modulating the proportion of these species suggests additional factors present in MSe involved in their regulation.

**Figure 5.**
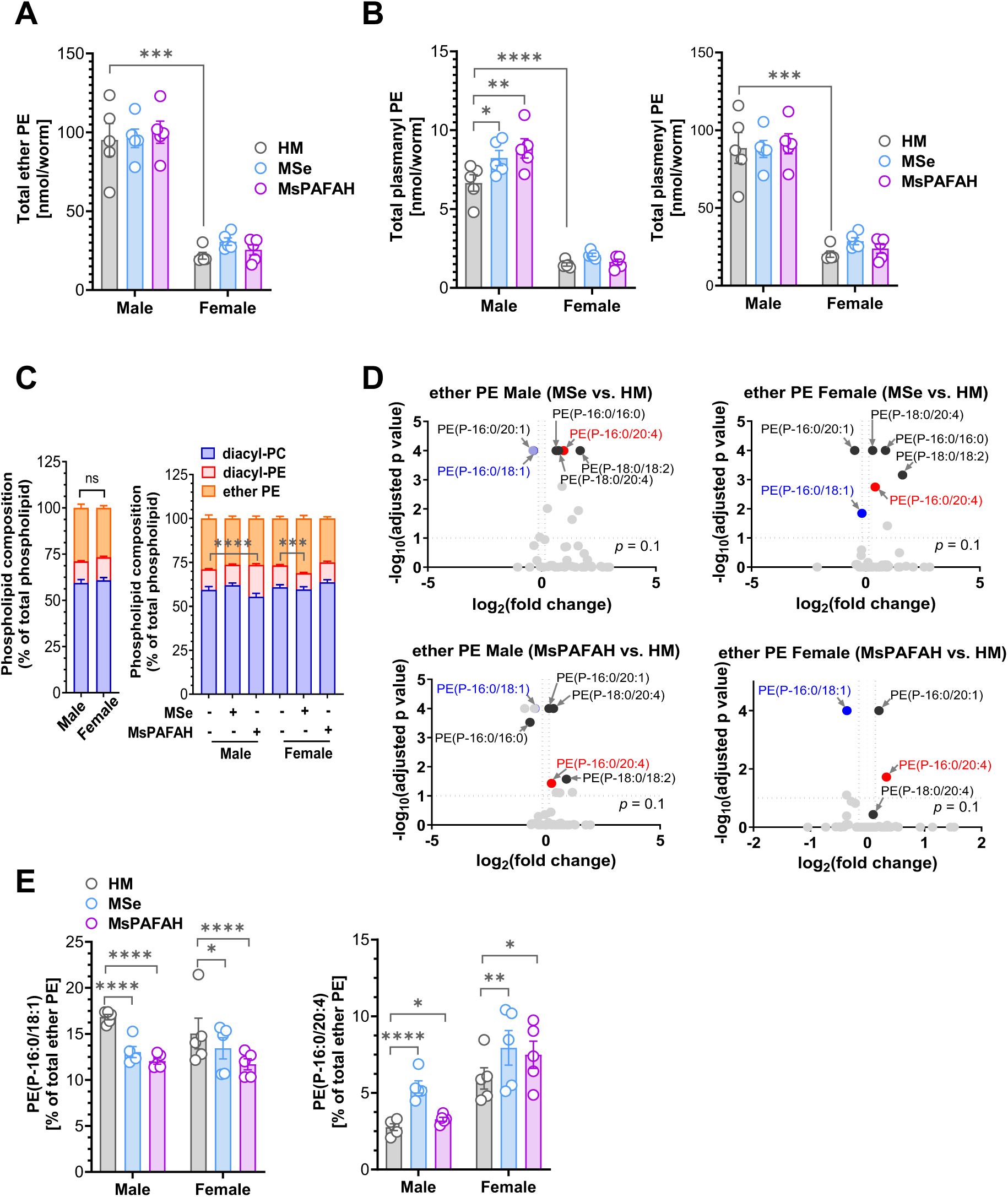
MsPAFAH remodels schistosomal ether PE abundance and composition in a sex-dependent manner. (A) UPLC-MS/MS analysis of the absolute amount (nmol/worm) of total ether phosphatidylethanolamine (PE) in male and female *ex-vivo* adult worms following MsPAFAH and MSe treatment. (B) UPLC-MS/MS analysis of the absolute amount (nmol/worm) of total plasmanyl PE (left panel) and plasmenyl PE (right panel) in male and female adult worms following MsPAFAH and Mse treatment. (C) Total diacyl-PC, diacyl-PE, and ether PE composition (% of total phospholipids) in male and female worms (left panel) and upon MsPAFAH and MSe treatment (right panel). (D) Volcano plots showing the differentially regulated ether PE species (relative intensities, % of total ether PE) upon MSe and MsPAFAH treatment in male and female worms. (E) Proportion of individual ether PE species (% of total ether PE). Proportions of the downregulated PE(P-16:0/18:1) and upregulated PE(P-16:0/20:4) species following MsPAFAH and MSe treatment. Data information and statistics: Results are representative of at least three independent experiments (5 worms in each group) and are expressed as means ± SEM. Statistical P values were calculated by ordinary two-way ANOVA + Dunnett’s post hoc test (A, B, left panel of C, D, E) or unpaired two-tailed *t*-test (right panel of C) *P < 0.05; **P < 0.01; ***P < 0.001; ****P < 0.0001; ns: not significant. For panel D, fold changes were calculated from the relative intensities (% of total PC or PE) of the species from MsPAFAH- or MSe-treated against HM-treated worms. The differentially regulated species were marked when the following criteria were met: 1) the proportion of the species (% of total ether PE) > 0.5%, 2) fold changes against HM control ≤ 0.9 or ≥ 1.1, 3) P values < 0.1 adjusted.

Together, exposure to MsPAFAH, and to a lesser extent to MSe, alters the ether PL composition, with an increase in the total plasmanyl ether PE levels, preferentially in male *S. mansoni* worms, with commonalities between males and females in some unsaturated plasmenyl PE species.

### Supplementation of fatty acids and precursors depleted from phospholipids by PAFAH exposure attenuates PAFAH’s toxicity preferentially in female worms

Our current data revealed that MsPAFAH induces remarkable changes in the PE and PC composition of *S. mansoni*, which correlate positively with decreased worm viability. In particular, it significantly reduced the proportion of the dominant schistosomal PL species, PC(16:0_20:1) and PE(18:0_22:4), irrespective of sex. Schistosomes have lost the ability to synthesize fatty acids *de novo*, but they can incorporate and modify fatty acids by chain elongation to build complex lipids when supplied with dietary sources (25, 26). This led us to test whether supplying exogenous FFA to MSe- and MsPAFAH-treated male and female adult worms would prevent the decline in viability and inhibit parasite death (Fig. 6). To this end, we supplemented adult worms with the FFA C20:1/eicosenoic acid and 22:4/adrenic acid during treatment with MSe and MsPAFAH to compensate for the reduction in PC(16:0_20:1) and PE(18:0_22:4), respectively (Fig. 6A). As expected, the supplementation of these FAs significantly increased the content of 20:1 and 22:4 in PE and PC in both male (Fig. B) and female (Fig. 6C) worms and compensate MSe- and MsPAFAH-induced disruption of these PE and PC species (Supplementary Fig. S5A-B). 18:0/stearic acid and DMPC (1,2-dimyristoyl-*sn*-glycero-phosphocholine) (major species not expected to be limiting) were supplemented as controls. Indeed, neither 18:0 stearic acid nor DMPC supplementation rescued worm viability after MSe and MsPAFAH treatment (Fig. 6D-E). However, kinetic studies revealed that supplementation with C20:1/eicosenoic acid tended to impair and delay the MSe/MsPAFAH-induced decrease in viability, especially in male worms (Fig. 6F). More strikingly, supplementation with 22:4/adrenic acid greatly delayed the MSe- and MsPAFAH-induced killing in males and largely prevented the decrease of viability in females up to 72 h (Fig. 6G). Of note, none of the supplemented lipids interfered with the viability of untreated worms (Fig. 6D-G).

**Figure 6.**
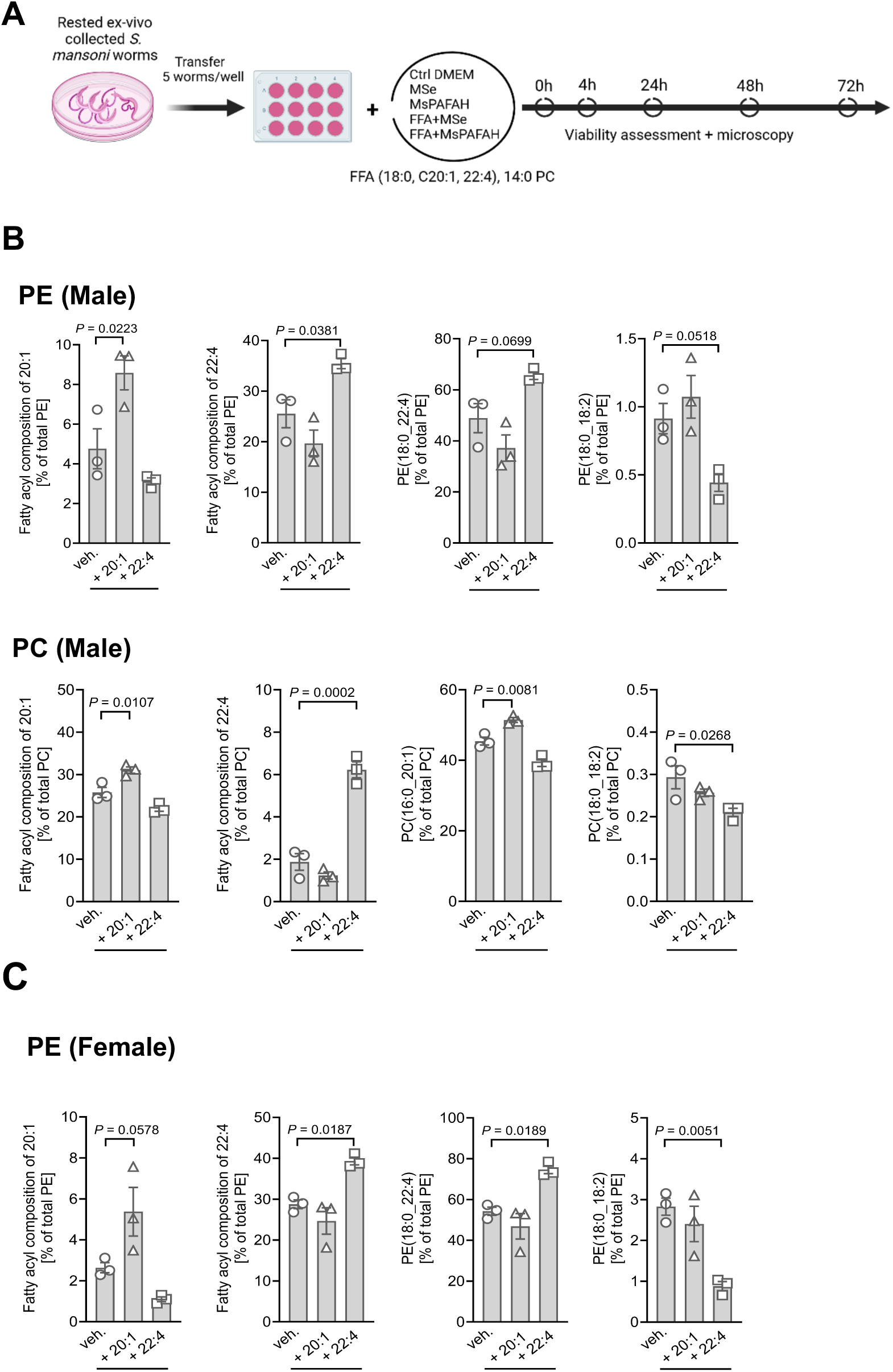
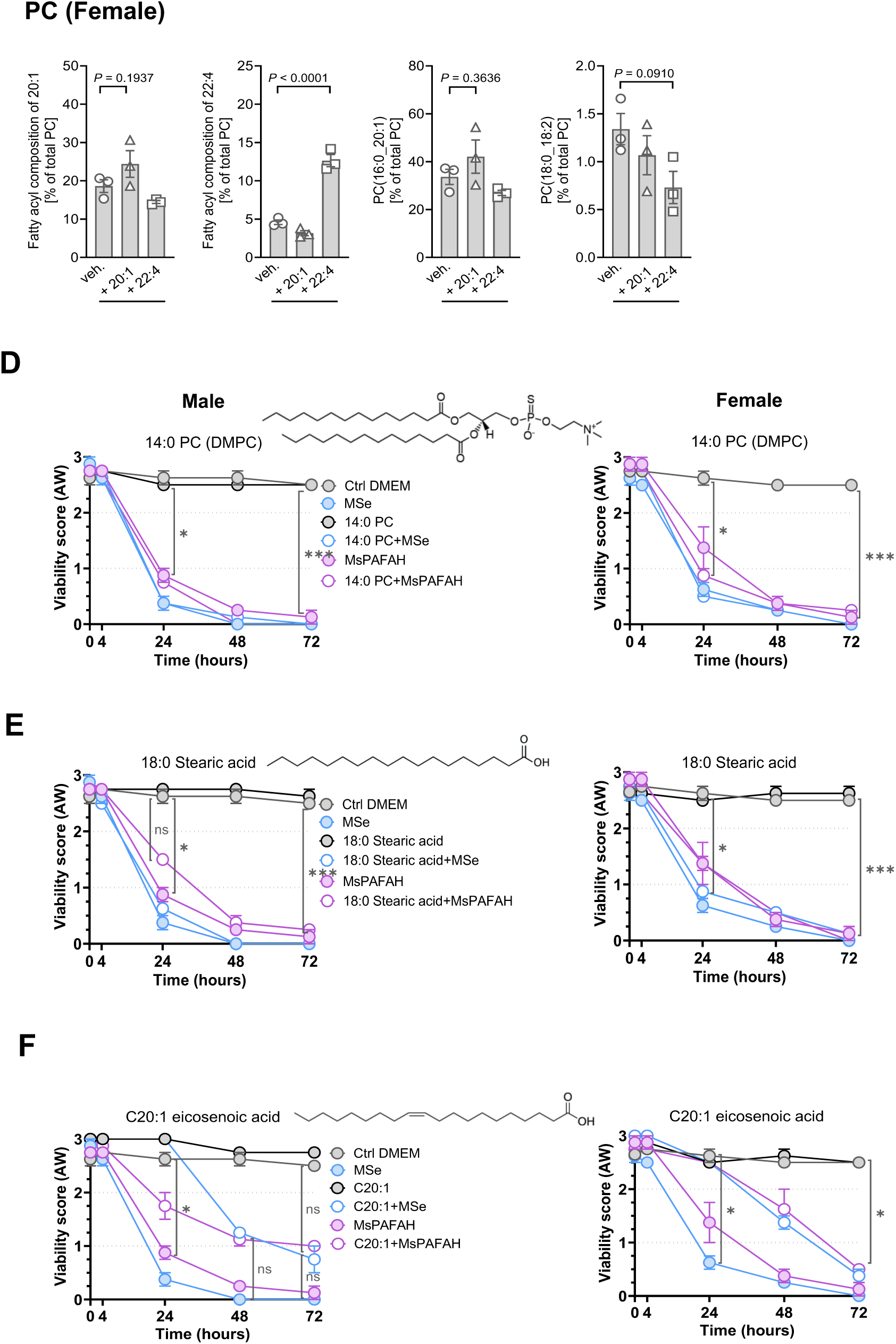
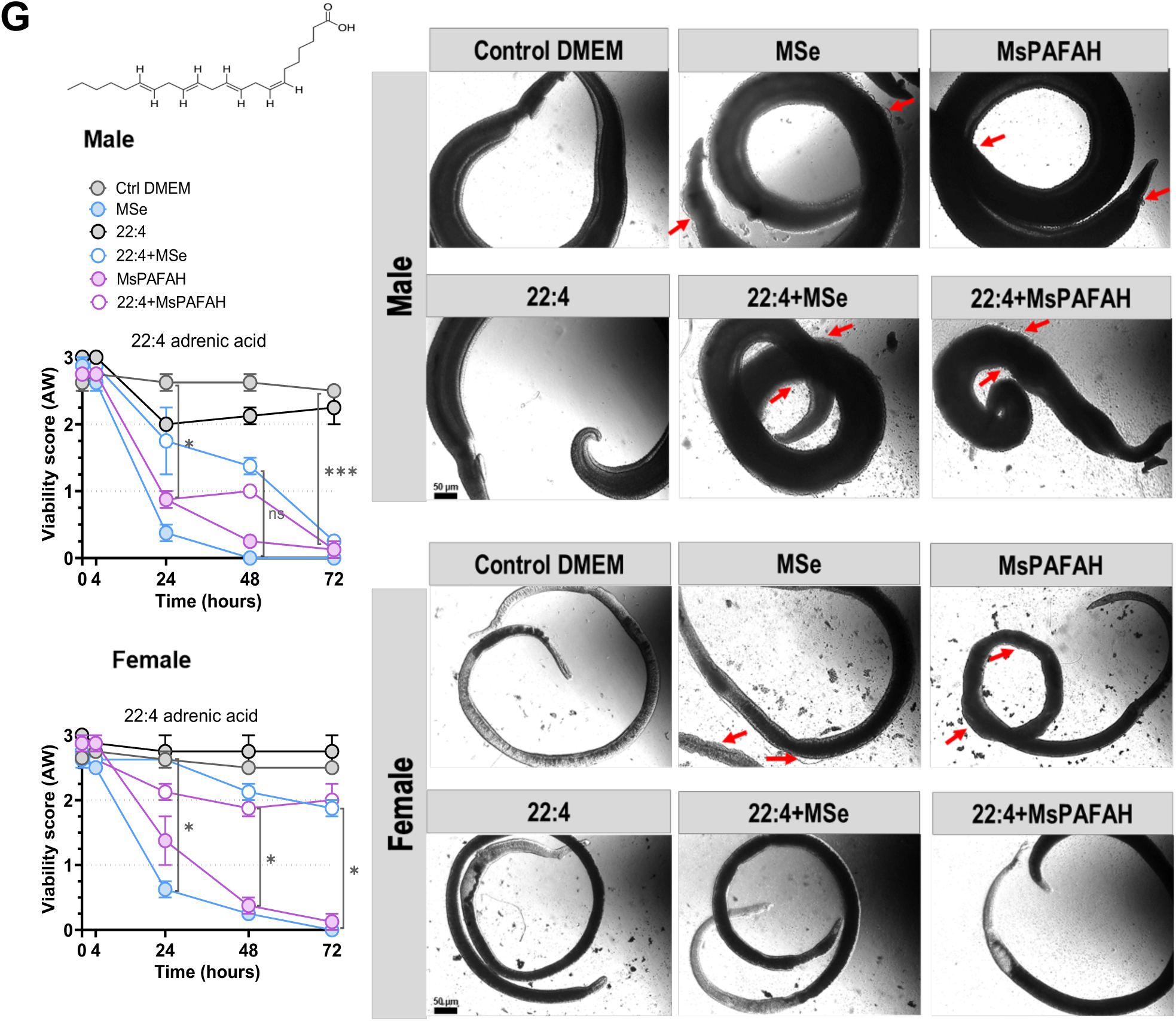
Supplementation of fatty acids and precursors depleted from phospholipids by PAFAH exposure attenuates PAFAH toxicity preferentially in female worms. (A) Experimental design of fatty acid supplementation and viability testing. (B) – (C) Respective diacyl-PE (upper panels) and diacyl-PC (lower panels) composition following fatty acid and precursors 20:1 and 22:4 supplementation to ex vivo recovered male (B) and female (C) adult worms in comparison to DMEM control (veh.) (D) - (G) Distinct fatty acid supplementation to *ex vivo* recovered male (left panel) and female (right panel) adult worms alone and in combination with MsPAFAH or MSe in comparison to DMEM control and viability assessment over time. A representative picture of each condition is shown in G for the supplementation of 22:4 adrenic acid. Red arrows show the effect of MsPAFAH and MSe, and in combination with 22:4 adrenic acid, on males and females. Data information and statistics: Results are representative of at least two to three independent experiments (5 worms per condition) and are expressed as means ± SEM. Asterisks show significant statistical differences analyzed using ordinary one-way ANOVA + Dunnett’s post hoc test (B - C) and two-way ANOVA followed by Tukey’s multiple comparison test (D-G). *P < 0.05; **P < 0.01; ***P < 0.001; ****P < 0.0001.

Microscopically, male and female parasites incubated in the presence of MSe and MsPAFAH alone appeared contracted and heavily granulated with disrupted tegument after 72 h incubation (Fig. 6G, right panel). Additionally, the worm tegumental tissue structure was remarkably affected showing bubble formation on the tegumental surface (red arrows). Unlike in male worms, the addition of 22:4/adrenic acid into the culture system abrogated the bubble formation and granulation in female worms, rendering their appearance comparable to untreated or 22:4 only treated worms.

Our data indicate that MsPAFAH-induced toxicity in *S. mansoni* is attenuated by supplying fatty acids, i.e. 22:4/adrenic acid and to a lesser extent C20:1/eicosenoic acid, which are crucial for the biosynthesis of membrane PLs that are preferentially depleted by MsPAFAH treatment. These effects were most evident at 24 to 48 h and less at 72 h, except for 22:4/adrenic acid in female worms, where consistent protection against worm death was observed up to 72 h.

Together, these results provide strong evidence for the critical role of PAFAH present in mouse serum as a potent schistosomal multi-stage agent in controlling the maintenance of tissue integrity and survival of *S. mansoni* by targeting schistosome body and tegumental PE and PC, specifically 20:1- and 22:4-containing species.

## Discussion

Regulated host-parasite interaction is a prerequisite for the survival of both the host and the parasite. Parasites, in particular, rely on specific adaptations to the host and the anatomic niche in which they reside. In the case of schistosomes, it is the blood system. Therefore, understanding the underlying developmental prerequisites for survival, such as interaction with soluble serum factors, opens avenues for better understanding host-specificity. This offers promise to identify suitable targets and schistomicidal mechanisms of host factors and molecules. In the case of schistosomes, mice can be a naturally occurring reservoir host despite being widely used as an experimental, preclinical model (10, 27, 62). We describe here, the identification of a soluble serum factor from mice, the platelet-activating factor acetylhydrolase (PAFAH), which is a bioactive enzyme and phospholipase present in mouse serum, as a highly potent schistosomicidal factor. We also identified other molecules (e.g. MUP10, adiponectin) which had schistosomicidal effects, however to a lesser extent and at higher concentrations, which could also contribute to the overall very strong killing effect of MSe. However, we focused in this study on deciphering the mode-of-action of PAFAH since it demonstrated the strongest schistosomicidal *in vitro* and *ex vivo* activity across all parasite developmental stages present in the definite host. We indeed found that PAFAH dysregulated specific parasite PLs, particularly certain species thereof (PE, PC) which overall demonstrates that the PL metabolism, which is central to maintaining tegument integrity and is thus one of the parasites’ Achilles heels, could open an avenue for pharmacological targeting in the future. While our findings underscore the significant role of MsPAFAH in regulating PL remodeling/metabolism, critical for the survival of *S. mansoni*, the current study predominantly relies on *in vitro* and *ex vivo* methodologies, which, while robust and controlled, may not fully capture the complexity of host-parasite interactions in *in vivo* settings. Translating these findings into therapeutic applications warrants further investigation.

Until now, investigations into targeting the parasite lipid metabolism for drug development are scarce. While previous studies provided valuable insights into the broader role of phospholipases in helminth biology and immune evasion, they failed to address how helminths, especially schistosomes, may interact with host lipid metabolism-related enzymes, such as PAFAH, during infection (26, 51, 55, 63, 64). Our findings reveal an increase of PAFAH, upon schistosome infection, in mouse serum, which we previously demonstrated to harbor soluble schistosomicidal factors (10), suggesting a key role of MsPAFAH in host-parasite interactions and the establishment and progression of *S. mansoni* infection in mice. In general, PAFAH is known to hydrolyze sn-2-acyl-bioactive lipids, especially the platelet-activating factor (PAF) (57, 61), and has previously garnered attention for its diverse physiological functions, including the modulation of immune responses and inflammatory processes in the host (11, 23, 57, 65–67). Interestingly, we noted a discrepancy in the impact of the anti-schistosomal capacity of PAFAH on the development and survival of schistosomes depending on the species of origin. Indeed, while MsPAFAH had detrimental effects on the survival and development of *S. mansoni ex vivo*, HuPAFAH instead promoted parasite survival and, subsequently also, development, as demonstrated previously (10, 27). Indeed, host PAFAH activity is closely related to its interactions with soluble lipoproteins, with high-density lipoprotein (HDL) mainly favoring a higher activity than low-density lipoprotein (LDL) (57, 68). In humans, 70% of plasma PAFAH is associated with LDL and only 30% with HDL (57, 68). In contrast, up to 70% of plasma PAFAH in mice is bound to HDL and expresses higher protein activity levels than species associated with LDL and low HDL particles (57, 68). Thus, the activity and target specificity of MsPAFAH are almost nine times higher than that of the human enzyme (69, 70). In addition, structural modeling of PAFAH from both species revealed distinct differences that could explain substrate specificity and increased activity. The comparison of the HuPAFAH structure with the AlphaFold2-derived model of MsPAFAH together with the sequence alignment clearly showed that while the catalytic triad is perfectly conserved and the HDL binding site is very similar in both orthologs, there are marked differences in the LDL binding site. Specifically, the triad Trp115/Leu116/Tyr205 is involved in LDL binding in HuPAFAH, but only Tyr205 is present in MsPAFAH, potentially explaining why MsPAFAH is mainly associated with HDL in mice.

Nevertheless, although the schistomicidal effects of MSe and MsPAFAH were similar in most cases, we sometimes observed a more potent killing effect of MSe, suggesting the potential presence of additional factors in serum with added effects. Also, *in vivo*, there might be endogenous inhibitors of PAFAH that are not present in the *in vitro* assay. Thus, eventually, the effects of MSe and MsPAFAH killing *in vitro* might be distinct in *in vivo* settings and need further investigation. Furthermore, the recombinant protein we used in our assays might be less potent or active when compared to the naturally occurring one despite clearly demonstrating specific enzymatic activity. This might result from improper folding or oxidative damage during recombinant expression or lack of post-translational modifications, and eventually due to the lack of HDL previously bound to the parasite tegument.

PAFAH-mediated damage was evident in the tegument and the schistosomes’ gut and reproductive organs, as revealed by CLSM and SEM, leading to an additional gross reduction in parasite reproductivity. Similar post-treatment phenotypes were previously reported for arylmethylamino steroids, a novel compound class affecting worm fitness, reproduction, and tissue morphology, including the gonads, tegument and gut (71). Although this compound’s target(s) are unknown, tegumental membrane destabilizing effects were discussed. The tegument of schistosomes, enriched in parasite-specific PL species, is a dynamic host-adapted interface between the parasite and its vascular environment. PLs as potential PAFAH targets are essential for schistosome survival and are increasingly recognized as mediators, or precursors thereof, in signal transduction and immune response modulation (26, 50). They constitute significant components of the layers that cover and protect schistosomes from diverse host factors, such as serum components (26, 59). Thus, tegument-specific PLs play a substantial role in host-parasite interactions, as evidenced by the roles of lyso-PC and lyso-phosphatidylserine in schistosomal host immune evasion (72, 73). These are also crucial for various pathophysiological processes related to cell survival and death of eukaryotes in general; for example, 1,2-dioleoyl-*sn*-glycero-phosphoinositol (PI(18:1/18:1)) in stress adaption (60), oxidized PLs in apoptosis and ferroptosis (74), and lipid mediators in inflammation (75), among others. We thus investigated whether PAFAH targeted specific PL species for hydrolysis, disrupting tegumental membrane integrity and allowing the influx of damaging ions and other molecules, eventually activating further internal signaling.

Notably, treatment with MsPAFAH, similarly to MSe, elevated total PC and PE levels, especially in male *S. mansoni*. This suggests a compensation mechanism in response to the MsPAFAH-mediated hydrolysis of lipids, which might correlate to schistosomes’ reported rapid membrane renewal capabilities (49). This renewal of the membrane complex is closely associated with increased schistosome lipid metabolism and related enzymes following stress events (76). These enzymes (e.g. acyltransferases present in schistosome) may be used by the worms to incorporate MsPAFAH-hydrolysis products (e.g. FFA and lyso-PL) into various PL species (77), such as PC and PE, thus compensating for membrane PL loss following MsPAFAH-dependent degradation, an effect we observed when we supplemented FFA to MSe or MsPAFAH treated schistosomes. Other studies suggested that the tegument is constantly renewed, either by acylation and deacylation of specific lipids directly in the tegument, and that the male worms shed their outer layer of the tegument, enriched in PLs, to potentially get rid of antibodies and attacking immune cells, resulting yet again in the upregulation of PL synthesis and levels (15, 49), especially in male worms. In line with our observations, previous electron microscopic investigations have identified LDL, but also HDL, to be attached to their receptors, the glycosylphosphatidylinositol (GPI)-linked low-molecular-weight proteins, on the outer layer of the tegument and dorsal regions of *ex vivo* flushed adult worms (78). Thus, HDL/LDL-bound PAFAH would potentially attach and hydrolyze tegument PLs, producing *S. mansoni* specific fatty acids (FA) and/or LysoPLs. In addition, bound HDL and LDL activate the cytons to produce membranous vacuoles to replace the missing tegument. The continuous tegumental shedding may disrupt the inner tegument layer and eventually lead to parasite death, especially in male schistosomes, possibly further explained by the fact that male schistosomes preferentially attach LDL and HDL particles to their tegument (78). Indeed, as previously reported, such sex-dependent differences with more effect on male worms were also observed along with higher PL content in males compared to females (50, 78). Whether the higher susceptibility of male worms to treatment is functionally linked to the higher content of total PLs or related to differences in the FA composition currently remains elusive.

Importantly, it was observed that specific PLs, such as ether PLs, with FA specific to schistosomes (e.g. 20:1, 18:0, 18:1(5Z)) are enriched in the tegument and determine the parasite susceptibility to stress and death signals (15, 49). These schistosome-specific FA and PLs might be used as feedback for the underlying tegumental cells regarding the integrity of the tegument (15, 49). To identify parasite-specific PLs in the tegument, whose levels are modulated by MsPAFAH, we ran a comprehensive and quantitative metabololipidomic analysis of specifically affected PC and PE. This revealed substantial changes in the PL profile (> 100 species), in particular of the dominant species PC(16:0_20:1) and PE (18:0_22:4), whose proportion was reduced in both male and female worms after MsPAFAH treatment, and, as such, might potentially represent specific (direct or indirect) targets of MsPAFAH. Interestingly, both MsPAFAH and MSe increased the proportion of PC(18:0_18:2) and PE(18:0_18:2) along with other diacyl- and ether-PC and -PE species in male and female worms. This implies several complex defense mechanisms activated in schistosomes by MsPAFAH treatment, damaging the worms’ tegument, gut, and reproductive organs. Remarkably, supplementation with the respective FFAs depleted in membrane PLs upon treatment with MsPAFAH almost entirely prevented the MsPAFAH-induced decrease in female viability while only partially in males. This is an essential finding with potential *in vivo* relevance concerning schistosome fecundity, reproductivity, and associated pathology. Moreover, paired female schistosomes have been demonstrated to rapidly incorporate, oxidize, modify, and turn over FAs, essential for egg production and integrity (79, 80).

## Conclusions

We provide here new insight into the critical and underresearched area of parasite-host specificity, deepening our understanding of the underlying factors of host-parasite interaction. To capitalize on this discovery for developing novel therapeutics against schistosomiasis, future studies should prioritize evaluating the efficacy of MsPAFAH or ideally analalogues thereof in *in vivo* models that mimic the dynamic biological environment of mammalian hosts. Further characterization of the functional PAFAH-induced changes in the lipid composition would unravel the mechanisms that control candidate bioactive lipids. This will allow us to understand the mode of action of PAFAH in more detail, address fundamental aspects of schistosome biology, reproductive development, and survival, and potentially identify accessible and druggable targets.

## Acknowledgements

We want to thank Julian Eifler, Ulla Henn, and Stephanie Fetzer for excellent technical help and maintenance of the schistosome life cycle. We gratefully acknowledge the technical support by Sabine Agel, Imaging Unit of the Biomedical Research Center Seltersberg (JLU Giessen), for conducting SEM analysis. Many thanks to the NIAID Schistosomiasis Resource Center of the Biomedical Research Institute (Rockville, MD). The NIAID Schistosomiasis Resource Center of the Biomedical Research Institute (Rockville, MD) provided the reagent through NIH-NIAID Contract HHSN272201700014I. NIH: *Biomphalaria glabrata* (NMRI) exposed to *Schistosoma mansoni* (NMRI).

## Author contributions

Conceptualization: CPdC, UFP, A

Data curation: EJ, UFP, ZR, A, AK, YH, SS, PB, MH, CGG, SH, FHF, CPdC

Formal analysis: EJ, UFP, ZR, A, AK, YH, SS, PB, MH, SH, FHF, CPdC

Funding acquisition: CPdC, A, UFP, AK, CGG, SH, FHF

Investigation: EJ, UFP, ZR, A, AK, YH, SS, PB, MH, JS, CGG, SH, FHF, CPdC

Methodology: EJ, UFP, ZR, A, AK, YH, SS, PB, MH, CGG, SH, FHF, CPdC

Project administration: CPdC, UFP, A, AK, SS, SH

Resources: AK, SS, MH, CGG, SH, FHF, CPdC

Software: SS, MH, SH, FHF Supervision: CPdC, UFP, A

Validation: EJ, UFP, ZR, A, AK, YH, SS, PB, MH, JS, CGG, SH, FHF, CPdC

Visualization: EJ, UFP, ZR, A, AK, YH, SS, PB, MH, JS, CGG, SH, FHF, CPdC

Writing – original draft: UFP, EJ, CPdC

Writing – review & editing: EJ, UFP, ZR, A, AK, YH, SS, PB, MH, JS, CGG, SH, FHF, CPdC

All authors reviewed the manuscript and approved the submitted version.

## Availability of data and materials

All data generated or analysed during this study are included in this published article and its supplementary information files.

## Ethics approval and consent to participate

The animal and mouse serum investigations were approved by the Bezirksregierung Oberbayern (license number AZ 55.2-1-54-2532-145-17). Investigations using human serum were approved by the local ethical committee of the Technical University of Munich (TUM) (Reference: 215/18S), and all individuals, included in the study, consented enrollment.

## Consent for publication

Not applicable.

## Supplementary Information

**Supplementary Figure S1.**
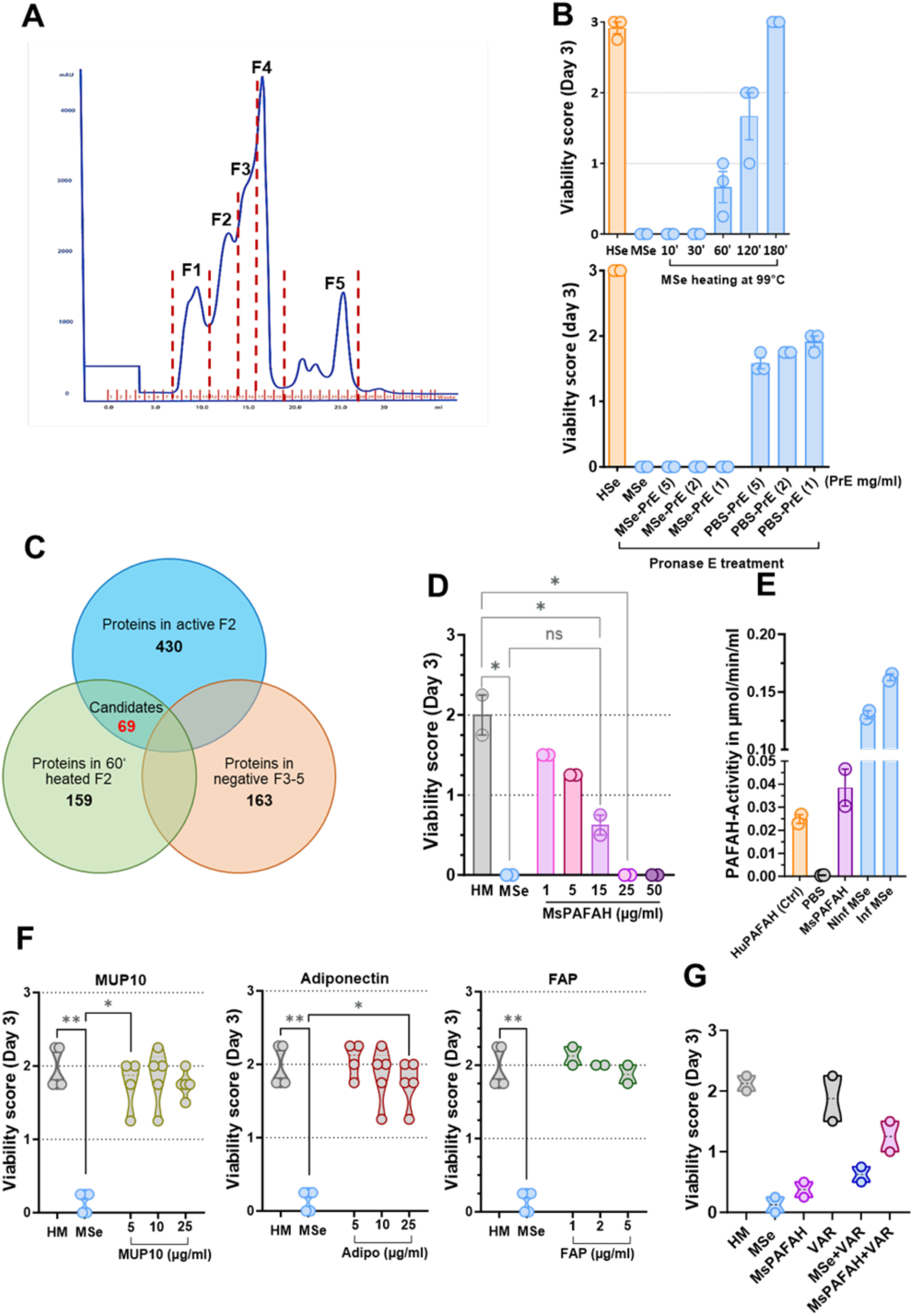
Large-scale fractionation of mouse serum using Supradex 200 size exclusion column and candidate molecule identification. (A) Mouse serum fractionation and pooling of recovered fractions (F1-5). (B) Upper panel: heating of mouse serum (MSe) active fraction F2 at different time points and effect on NTS viability at day 3. Lower panel: Pronase E (PrE)-digested mouse serum (MSe) and PBS (as control) and effect on NTS at day 3. (C) Venn diagram of refined proteins in active and inactive mouse serum fractions and heated fraction F2 with the candidate molecules. (D) Concentration-dependent effect of MsPAFAH as compared to MSe and HM control on NTS at day 3. (E) Enzymatic activity of recombinant MsPAFAH (44 µg/ml) as compared to infected (Inf) and non-infected (Ninf) MSe, and to positive control (HuPAFAH) and PBS. (F) Concentration-dependent effect MUP10, Adiponectin, and FAP as compared to Mse and HM control on NTS at day 3. (G) Effect of MSe and MsPAFAH on NTS viability in the presence or absence of the phospholipase A2 selective inhibitor varespladib (VAR) at day 3.

**Supplementary Figure S2.**
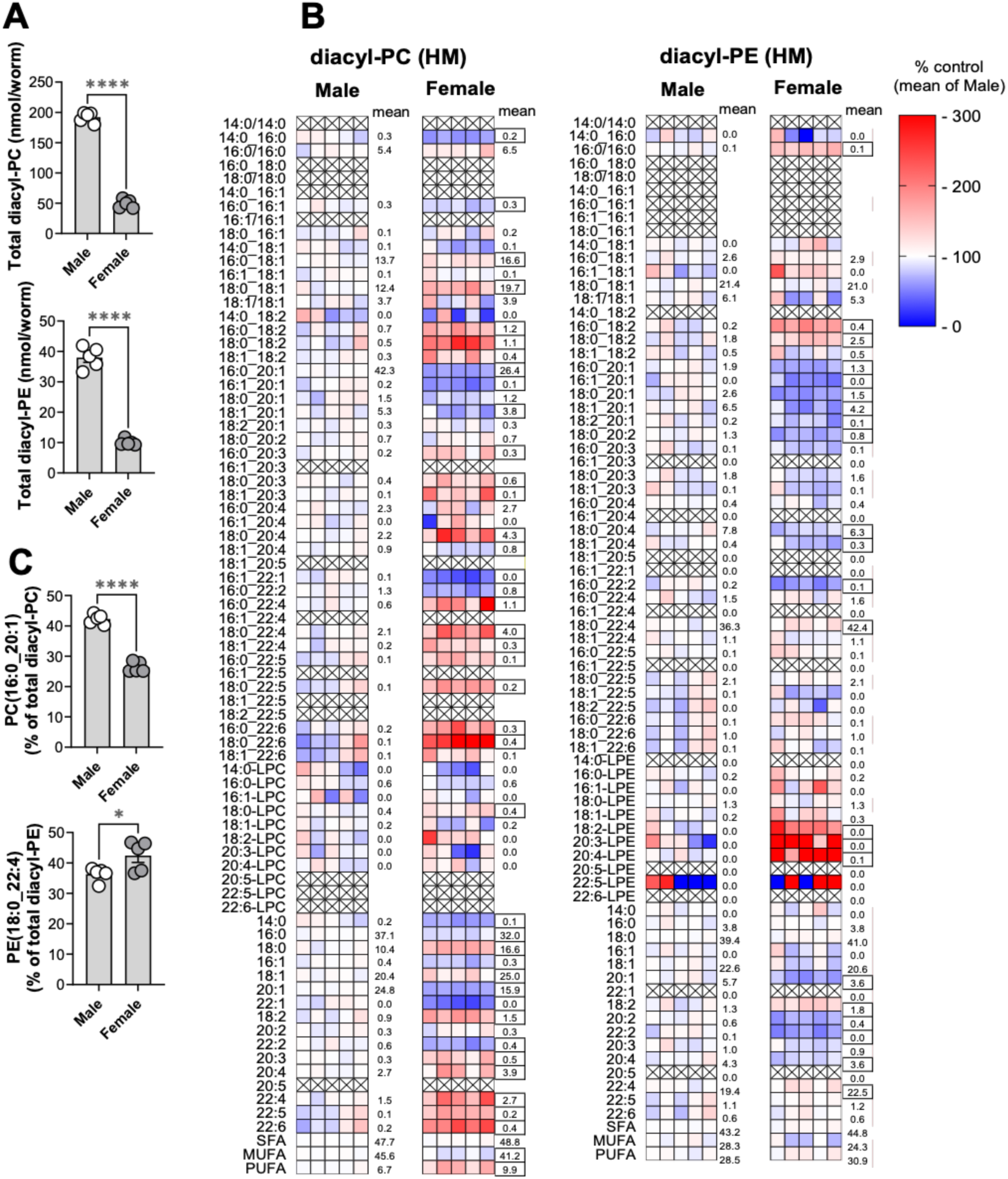
Gender differences of tegument diacyl-PC and -PE profiles in *S. mansoni* adult worms. Lipids were extracted from male and female S. mansoni worms, phosphatidylcholine (PC) and phosphatidylethanolamine (PE) were analyzed by UPLC-MS/MS. (A) Total diacyl-PC and -PE amount (pmol/worm). (B) The proportion of individual diacyl-PC,- PE species and fatty acid distribution (% of total diacyl-PC or -PE; SFA: saturated fatty acids, MUFA: monounsaturated fatty acids, PUFA: polyunsaturated fatty acids). The color code represents the fold change (% of the mean of male worms). The numbers indicate the mean values of the individual groups, and the differentially regulated species with P < 0.1 were marked by squares. (C) The proportion of the exemplary PC(16:0_20:1) and PE(18:0_22:4). Data are presented as A, C) mean ± S.E.M. or B) mean, n = 5 worms for each gender. *P < 0.05, ****P< 0.0001, two-tailed unpaired student t-test.

**Supplementary Figure S3.**
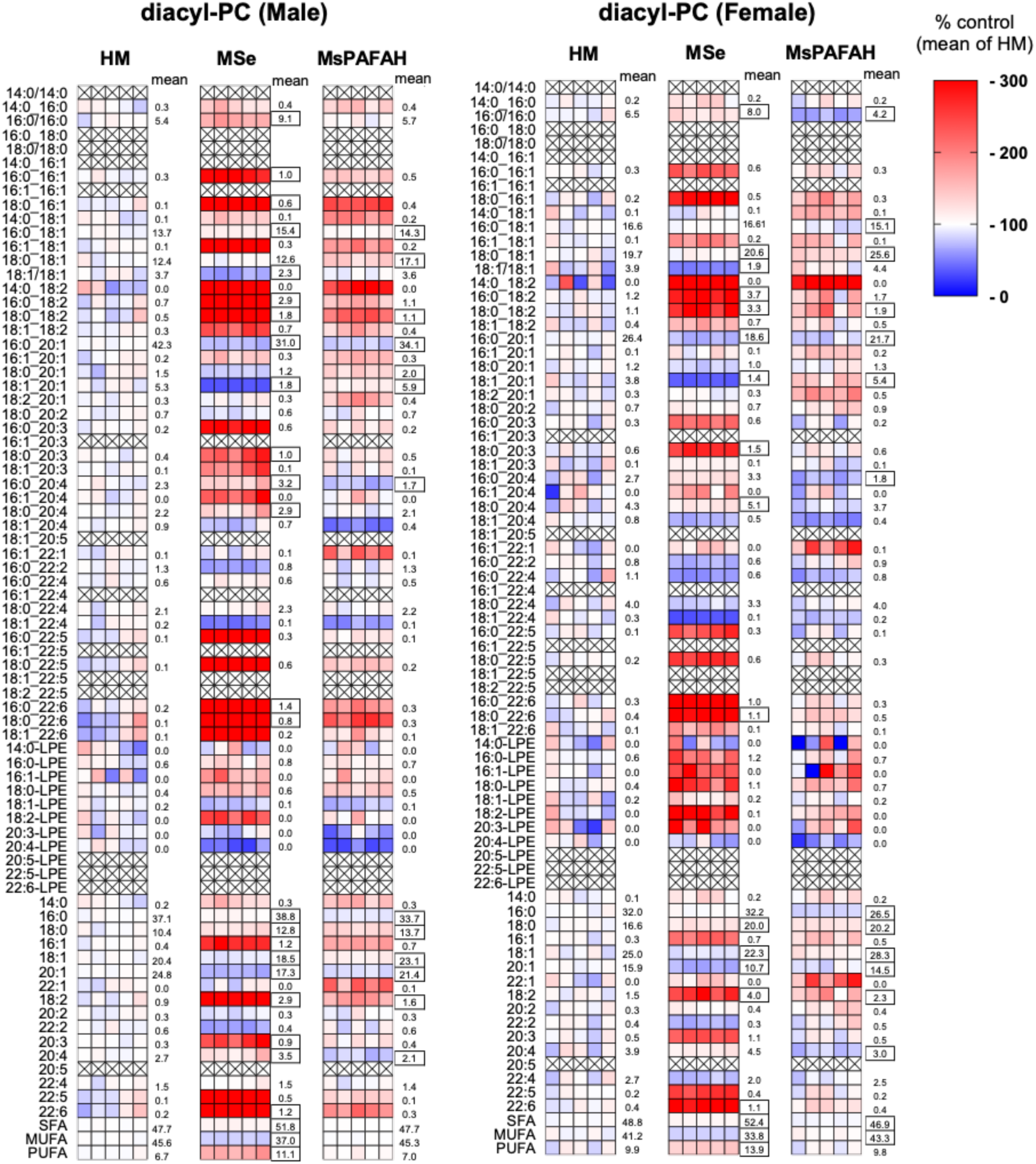
Diacyl-PC profile of the *S. mansoni* adult worms after MSe and MsPAFAH treatment. Lipids were extracted, phosphatidylcholine (PC) was analyzed by UPLC-MS/MS. The proportion of individual diacyl-PC (% of total diacyl-PC) and fatty acid distribution (% of diacyl-total PC; SFA: saturated fatty acids, MUFA: monounsaturated fatty acids, PUFA: polyunsaturated fatty acids). The color code represents the fold change (% of the mean of HM control). The numbers indicate the mean values of the individual groups and the differentially regulated species. Statistical P values were calculated by ordinary two-way ANOVA + Dunnett’s post hoc tests with P < 0.1 were marked.

**Supplementary Figure S4.**
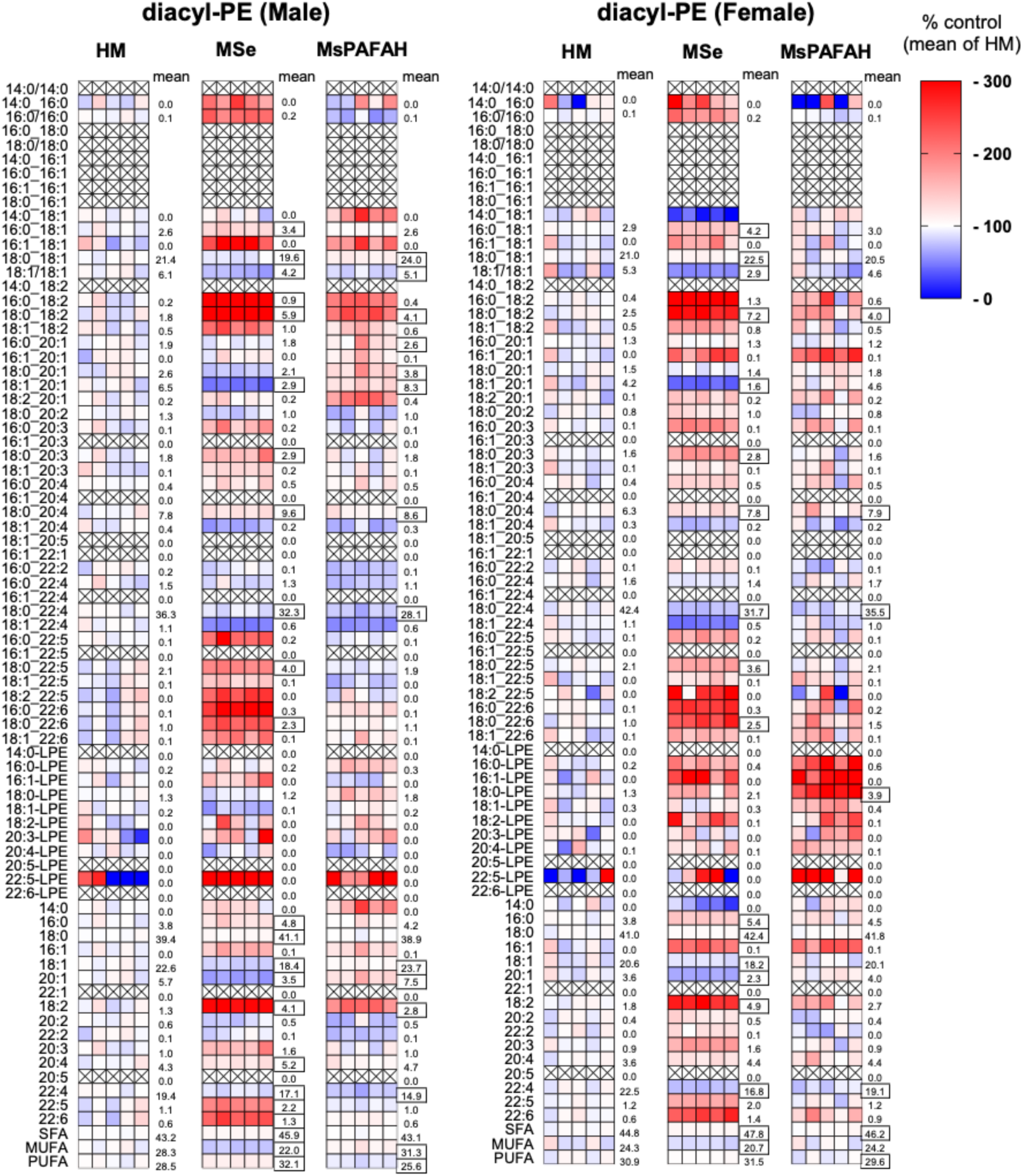
Diacyl-PE profile of the *S. mansoni* adult worms after MSe and MsPAFAH treatment. Lipids were extracted, phosphatidylethanolamine (PE) was analyzed by UPLC-MS/MS. The proportion of individual diacyl-PE (% of total diacyl-PE) and fatty acid distribution (% of total diacyl-PE; SFA: saturated fatty acids, MUFA: monounsaturated fatty acids, PUFA: polyunsaturated fatty acids). The color code represents the fold change (% of the mean of HM control). The numbers indicate the mean values of the individual groups and the differentially regulated species. Statistical P values were calculated by ordinary two-way ANOVA + Dunnett’s post hoc test with P < 0.1 were marked.

**Supplementary Figure S5.**
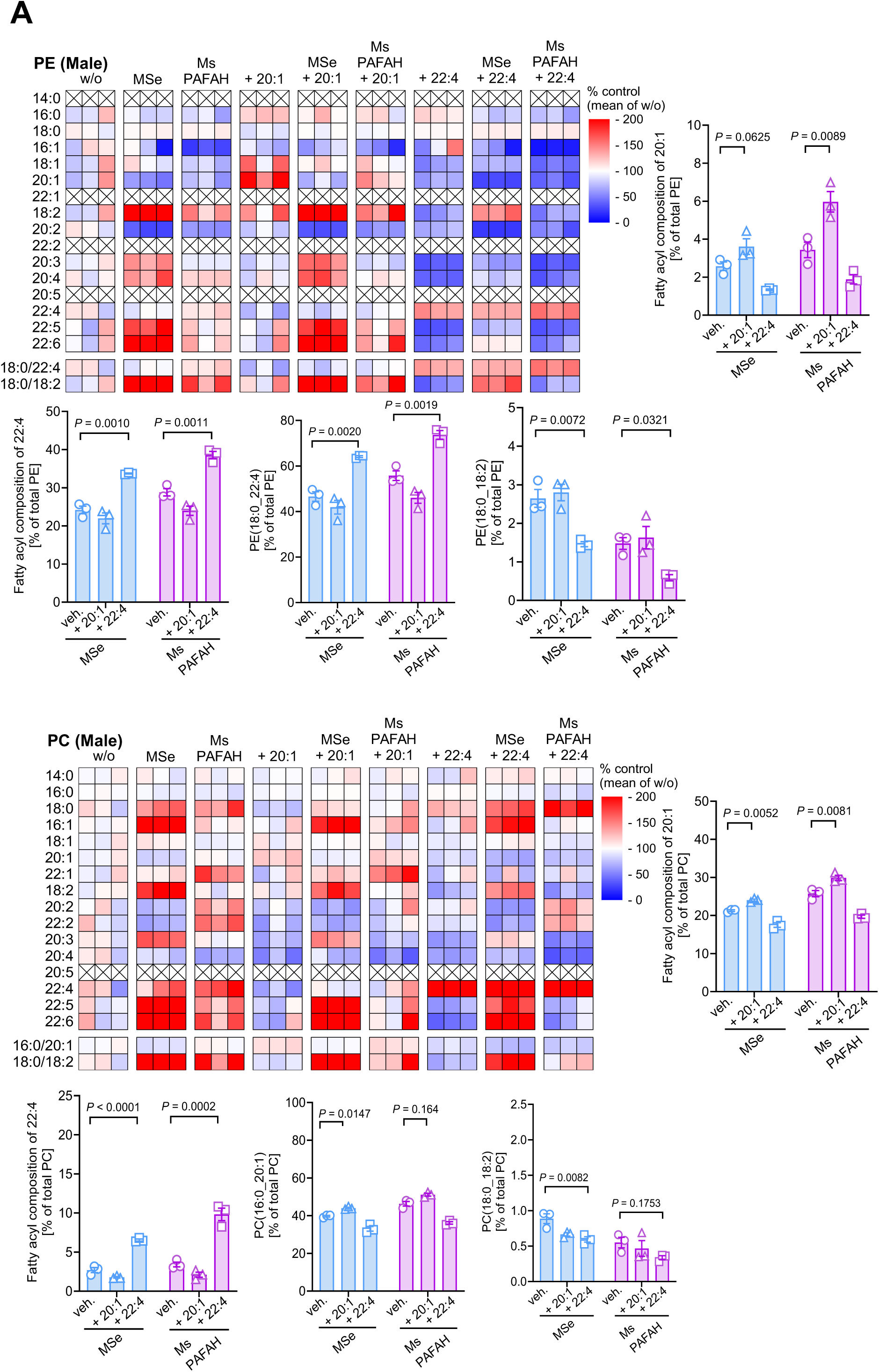
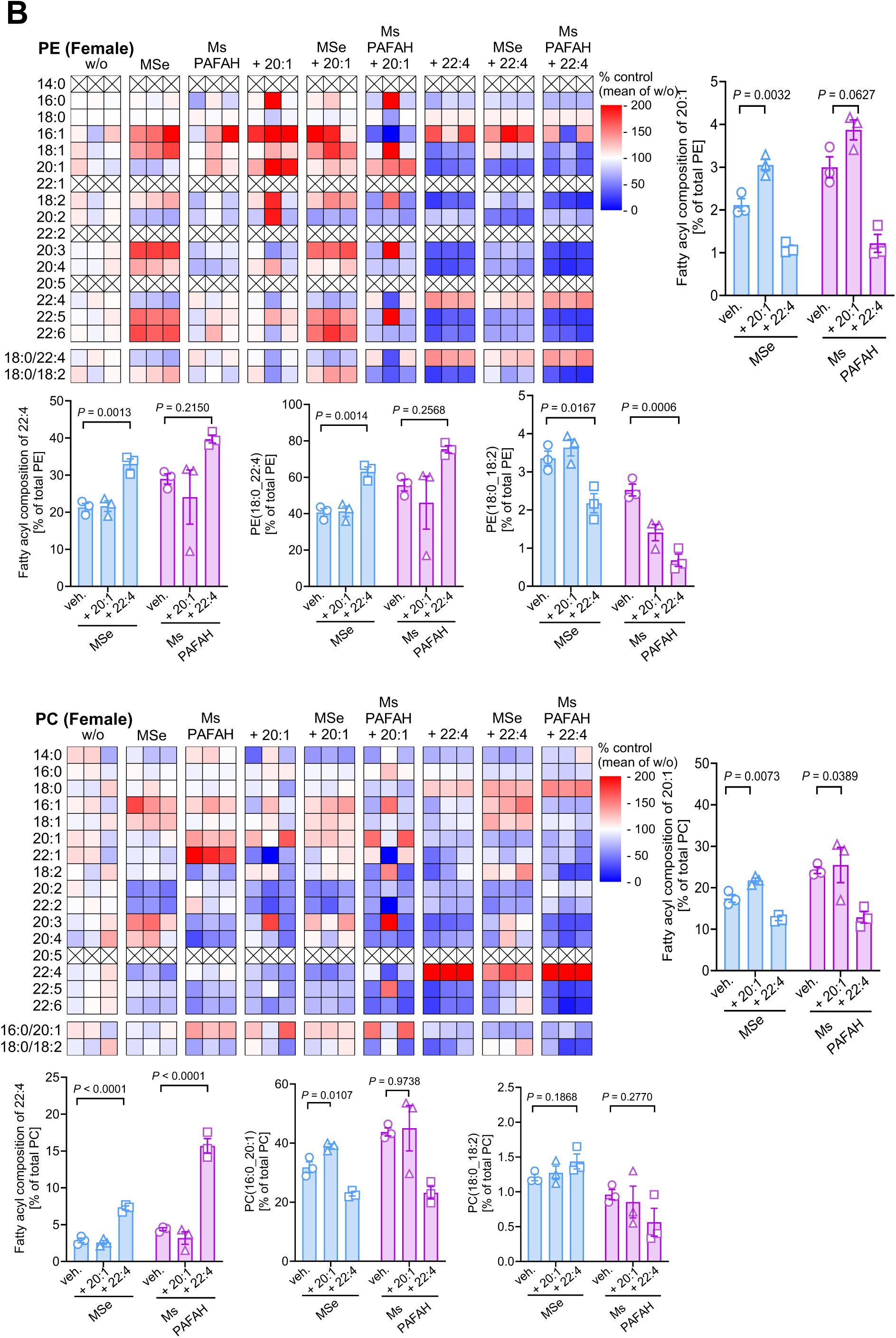
Supplementation of fatty acids and precursors increases the uptake of respective PE and PC species by male worms. (A-B) Respective diacyl-PE (upper panels) and diacyl-PC (lower panels) composition following fatty acid and precursors supplementation to *ex vivo* recovered male adult worms in combination with MsPAFAH or Mse in comparison to DMEM control (veh.) Results are representative of at least two-three independent experiments (5 worms per condition) and are expressed as means ± SEM. Asterisks show significant statistical differences analyzed using ordinary One-Way ANOVA + Dunnett’s post hoc test. *P < 0.05; **P < 0.01; ***P < 0.001; ****P < 0.0001

## References

1. McManus DP, Dunne DW, Sacko M, Utzinger J, Vennervald BJ, Zhou XN. Schistosomiasis. Nat Rev Dis Primers. 2018;4(1):13.

2. Tchuem Tchuente LA, Rollinson D, Stothard JR, Molyneux D. Moving from control to elimination of schistosomiasis in sub-Saharan Africa: time to change and adapt strategies. Infect Dis Poverty. 2017;6(1):42.

3. van der Werf MJ, de Vlas SJ, Brooker S, Looman CW, Nagelkerke NJ, Habbema JD, et al. Quantification of clinical morbidity associated with schistosome infection in sub-Saharan Africa. Acta Trop. 2003;86(2-3):125–39.

4. Colley DG, Bustinduy AL, Secor WE, King CH. Human schistosomiasis. Lancet. 2014;383(9936):2253–64.

5. Arsuaga M, Diaz-Menendez M, Gobbi FG. Autochthonous schistosomiasis in Europe: A silent threat. Travel Med Infect Dis. 2022;45:102244.

6. Rothe C, Zimmer T, Schunk M, Wallrauch C, Helfrich K, Gultekin F, et al. Developing Endemicity of Schistosomiasis, Corsica, France. Emerg Infect Dis. 2021;27(1):319–21.

7. Valentim CL, Cioli D, Chevalier FD, Cao X, Taylor AB, Holloway SP, et al. Genetic and molecular basis of drug resistance and species-specific drug action in schistosome parasites. Science. 2013;342(6164):1385-9.

8. Wang W, Wang L, Liang YS. Susceptibility or resistance of praziquantel in human schistosomiasis: a review. Parasitol Res. 2012;111(5):1871–7.

9. WHO guideline on control and elimination of human schistosomiasis. WHO Guidelines Approved by the Guidelines Review Committee. Geneva2022.

10. Anisuzzaman, Frahm S, Prodjinotho UF, Bhattacharjee S, Verschoor A, Prazeres da Costa C. Host-Specific Serum Factors Control the Development and Survival of Schistosoma mansoni. Frontiers in Immunology. 2021;12(1384).

11. Goncalves-Silva G, Vieira L, Cosenza-Contreras M, Souza AFP, Costa DC, Castro-Borges W. Profiling the serum proteome during Schistosoma mansoni infection in the BALB/c mice: A focus on the altered lipid metabolism as a key modulator of host-parasite interactions. Front Immunol. 2022;13:955049.

12. Wilson RA. The saga of schistosome migration and attrition. Parasitology. 2009;136(12):1581–92.

13. Maizels RM, Smits HH, McSorley HJ. Modulation of Host Immunity by Helminths: The Expanding Repertoire of Parasite Effector Molecules. Immunity. 2018;49(5):801–18.

14. Kardoush MI, Ward BJ, Ndao M. Identification of Candidate Serum Biomarkers for Schistosoma mansoni Infected Mice Using Multiple Proteomic Platforms. PLoS One. 2016;11(5):e0154465.

15. Van Hellemond JJ, Retra K, Brouwers JF, van Balkom BW, Yazdanbakhsh M, Shoemaker CB, et al. Functions of the tegument of schistosomes: clues from the proteome and lipidome. Int J Parasitol. 2006;36(6):691–9.

16. Holdgate GA, Meek TD, Grimley RL. Mechanistic enzymology in drug discovery: a fresh perspective. Nat Rev Drug Discov. 2018;17(1):78.

17. Wagner MP, Formaglio P, Gorgette O, Dziekan JM, Huon C, Berneburg I, et al. Human peroxiredoxin 6 is essential for malaria parasites and provides a host-based drug target. Cell Rep. 2022;39(11):110923.

18. Vanauberg D, Schulz C, Lefebvre T. Involvement of the pro-oncogenic enzyme fatty acid synthase in the hallmarks of cancer: a promising target in anti-cancer therapies. Oncogenesis. 2023;12(1):16.

19. Bayerl F, Meiser P, Donakonda S, Hirschberger A, Lacher SB, Pedde AM, et al. Tumor-derived prostaglandin E2 programs cDC1 dysfunction to impair intratumoral orchestration of anti-cancer T cell responses. Immunity. 2023;56(6):1341–58 e11.

20. Stine ZE, Schug ZT, Salvino JM, Dang CV. Targeting cancer metabolism in the era of precision oncology. Nat Rev Drug Discov. 2022;21(2):141–62.

21. Braschi S, Curwen RS, Ashton PD, Verjovski-Almeida S, Wilson A. The tegument surface membranes of the human blood parasite Schistosoma mansoni: a proteomic analysis after differential extraction. Proteomics. 2006;6(5):1471–82.

22. Leow CY, Willis C, Hofmann A, Jones MK. Structure-function analysis of apical membrane-associated molecules of the tegument of schistosome parasites of humans: prospects for identification of novel targets for parasite control. Br J Pharmacol. 2015;172(7):1653–63.

23. Espinoza B, Silman I, Arnon R, Tarrab-Hazdai R. Phosphatidylinositol-specific phospholipase C induces biosynthesis of acetylcholinesterase via diacylglycerol in Schistosoma mansoni. Eur J Biochem. 1991;195(3):863–70.

24. Giera M, Kaisar MMM, Derks RJE, Steenvoorden E, Kruize YCM, Hokke CH, et al. The Schistosoma mansoni lipidome: Leads for immunomodulation. Anal Chim Acta. 2018;1037:107–18.

25. Meyer F, Meyer H, Bueding E. Lipid metabolism in the parasitic and free-living flatworms, Schistosoma mansoni and Dugesia dorotocephala. Biochim Biophys Acta. 1970;210(2):257–66.

26. Retra K, deWalick S, Schmitz M, Yazdanbakhsh M, Tielens AG, Brouwers JF, et al. The tegumental surface membranes of Schistosoma mansoni are enriched in parasite-specific phospholipid species. Int J Parasitol. 2015;45(9-10):629–36.

27. Frahm S, Anisuzzaman A, Prodjinotho UF, Vejzagic N, Verschoor A, Prazeres da Costa C. A novel cell-free method to culture Schistosoma mansoni from cercariae to juvenile worm stages for in vitro drug testing. PLoS Negl Trop Dis. 2019;13(1):e0006590.

28. Vejzagic N, Prodjinotho UF, El-Khafif N, Huang R, Simeonov A, Spangenberg T, et al. Identification of hit compounds with anti-schistosomal activity on in vitro generated juvenile worms in cell-free medium. PLoS Negl Trop Dis. 2021;15(5):e0009432.

29. Basch PF. Cultivation of Schistosoma mansoni in vitro. I. Establishment of cultures from cercariae and development until pairing. J Parasitol. 1981;67(2):179–85.

30. Lacorcia M, Kugyelka R, Spechtenhauser L, Prodjinotho UF, Hamway Y, Spangenberg T, et al. Praziquantel Reduces Maternal Mortality and Offspring Morbidity by Enhancing Anti-Helminthic Immune Responses. Front Immunol. 2022;13:878029.

31. Clegg JA. IN VITRO CULTIVATION OF SCHISTOSOMA MANSONI. Exp Parasitol. 1965;16:133–47.

32. Misslin C, Velasco-Estevez M, Albert M, O’Sullivan SA, Dev KK. Phospholipase A2 is involved in galactosylsphingosine-induced astrocyte toxicity, neuronal damage and demyelination. PLoS One. 2017;12(11):e0187217.

33. Salvador GHM, Gomes AAS, Bryan-Quiros W, Fernandez J, Lewin MR, Gutierrez JM, et al. Structural basis for phospholipase A(2)-like toxin inhibition by the synthetic compound Varespladib (LY315920). Sci Rep. 2019;9(1):17203.

34. Hamway Y, Zimmermann K, Blommers MJJ, Sousa MV, Häberli C, Kulkarni S, et al. Modulation of Host-Parasite Interactions with Small Molecules Targeting Schistosoma mansoni microRNAs. ACS Infect Dis. 2022;8(10):2028–34.

35. Ramirez B, Bickle Q, Yousif F, Fakorede F, Mouries MA, Nwaka S. Schistosomes: challenges in compound screening. Expert Opin Drug Discov. 2007;2(s1):S53–61.

36. Lombardo FC, Pasche V, Panic G, Endriss Y, Keiser J. Life cycle maintenance and drug-sensitivity assays for early drug discovery in Schistosoma mansoni. Nat Protoc. 2019;14(2):461–81.

37. Neves RH, de Lamare Biolchini C, Machado-Silva JR, Carvalho JJ, Branquinho TB, Lenzi HL, et al. A new description of the reproductive system of Schistosoma mansoni (Trematoda: Schistosomatidae) analyzed by confocal laser scanning microscopy. Parasitol Res. 2005;95(1):43–9.

38. Beckmann S, Grevelding CG. Imatinib has a fatal impact on morphology, pairing stability and survival of adult Schistosoma mansoni in vitro. Int J Parasitol. 2010;40(5):521–6.

39. Mokosch AS, Gerbig S, Grevelding CG, Haeberlein S, Spengler B. High-resolution AP-SMALDI MSI as a tool for drug imaging in Schistosoma mansoni. Anal Bioanal Chem. 2021;413(10):2755–66.

40. Palmer A, Phapale P, Chernyavsky I, Lavigne R, Fay D, Tarasov A, et al. FDR-controlled metabolite annotation for high-resolution imaging mass spectrometry. Nat Methods. 2017;14(1):57–60.

41. Paschke C, Leisner A, Hester A, Maass K, Guenther S, Bouschen W, et al. Mirion--a software package for automatic processing of mass spectrometric images. J Am Soc Mass Spectrom. 2013;24(8):1296–306.

42. Koeberle A, Shindou H, Harayama T, Shimizu T. Role of lysophosphatidic acid acyltransferase 3 for the supply of highly polyunsaturated fatty acids in TM4 Sertoli cells. Faseb j. 2010;24(12):4929–38.

43. Thürmer M, Gollowitzer A, Pein H, Neukirch K, Gelmez E, Waltl L, et al. PI(18:1/18:1) is a SCD1-derived lipokine that limits stress signaling. Nat Commun. 2022;13(1):2982.

44. Haeberlein S, Angrisano A, Quack T, Lu Z, Kellershohn J, Blohm A, et al. Identification of a new panel of reference genes to study pairing-dependent gene expression in Schistosoma mansoni. Int J Parasitol. 2019;49(8):615–24.

45. Livak KJ, Schmittgen TD. Analysis of relative gene expression data using real-time quantitative PCR and the 2(-Delta Delta C(T)) Method. Methods. 2001;25(4):402–8.

46. Madeira F, Madhusoodanan N, Lee J, Eusebi A, Niewielska A, Tivey ARN, et al. The EMBL-EBI Job Dispatcher sequence analysis tools framework in 2024. Nucleic Acids Res. 2024.

47. Wilkins MR, Gasteiger E, Bairoch A, Sanchez JC, Williams KL, Appel RD, et al. Protein identification and analysis tools in the ExPASy server. Methods Mol Biol. 1999;112:531–52.

48. Waterhouse A, Bertoni M, Bienert S, Studer G, Tauriello G, Gumienny R, et al. SWISS-MODEL: homology modelling of protein structures and complexes. Nucleic Acids Res. 2018;46(W1):W296–W303.

49. Perez H, Terry RJ. The killing of adult Schistosoma mansoni in vitro in the presence of antisera to host antigenic determinants and peritoneal cells. Int J Parasitol. 1973;3(4):499–503.

50. Winkelmann F, Gesell Salazar M, Hentschker C, Michalik S, Machacek T, Scharf C, et al. Comparative proteome analysis of the tegument of male and female adult Schistosoma mansoni. Sci Rep. 2022;12(1):7569.

51. Angeles JMM, Mercado VJP, Rivera PT. Behind Enemy Lines: Immunomodulatory Armamentarium of the Schistosome Parasite. Front Immunol. 2020;11:1018.

52. Koopman JPR, Houlder EL, Janse JJ, Casacuberta-Partal M, Lamers OAC, Sijtsma JC, et al. Safety and infectivity of female cercariae in Schistosoma-naive, healthy participants: a controlled human Schistosoma mansoni infection study. EBioMedicine. 2023;97:104832.

53. Wilson RA, Barnes PE. The tegument of Schistosoma mansoni: observations on the formation, structure and composition of cytoplasmic inclusions in relation to tegument function. Parasitology. 1974;68(2):239–58.

54. Morris GP. Fine structure of the gut epithelium of Schistosoma mansoni. Experientia. 1968;24(5):480–2.

55. Du X, McManus DP, Fogarty CE, Jones MK, You H. Schistosoma mansoni Fibroblast Growth Factor Receptor A Orchestrates Multiple Functions in Schistosome Biology and in the Host-Parasite Interplay. Front Immunol. 2022;13:868077.

56. Kellershohn J, Thomas L, Hahnel SR, Grünweller A, Hartmann RK, Hardt M, et al. Insects in anthelminthics research: Lady beetle-derived harmonine affects survival, reproduction and stem cell proliferation of Schistosoma mansoni. PLoS Negl Trop Dis. 2019;13(3):e0007240.

57. Stafforini DM. Biology of platelet-activating factor acetylhydrolase (PAF-AH, lipoprotein associated phospholipase A2). Cardiovasc Drugs Ther. 2009;23(1):73–83.

58. Dong L, Li Y, Wu H. Platelet activating-factor acetylhydrolase II: A member of phospholipase A2 family that hydrolyzes oxidized phospholipids. Chem Phys Lipids. 2021;239:105103.

59. Brouwers JF, Van Hellemond JJ, van Golde LM, Tielens AG. Ether lipids and their possible physiological function in adult Schistosoma mansoni. Mol Biochem Parasitol. 1998;96(1-2):49–58.

60. Thurmer M, Gollowitzer A, Pein H, Neukirch K, Gelmez E, Waltl L, et al. PI(18:1/18:1) is a SCD1-derived lipokine that limits stress signaling. Nat Commun. 2022;13(1):2982.

61. Kriska T, Marathe GK, Schmidt JC, McIntyre TM, Girotti AW. Phospholipase action of platelet-activating factor acetylhydrolase, but not paraoxonase-1, on long fatty acyl chain phospholipid hydroperoxides. J Biol Chem. 2007;282(1):100–8.

62. Catalano S, Sene M, Diouf ND, Fall CB, Borlase A, Leger E, et al. Rodents as Natural Hosts of Zoonotic Schistosoma Species and Hybrids: An Epidemiological and Evolutionary Perspective From West Africa. J Infect Dis. 2018;218(3):429–33.

63. Du X, Jones MK, Nawaratna SSK, Ranasinghe S, Xiong C, Cai P, et al. Gene Expression in Developmental Stages of Schistosoma japonicum Provides Further Insight into the Importance of the Schistosome Insulin-Like Peptide. Int J Mol Sci. 2019;20(7).

64. Shen J, Zhao S, Peng M, Li Y, Zhang L, Li X, et al. Macrophage-mediated trogocytosis contributes to destroying human schistosomes in a non-susceptible rodent host, Microtus fortis. Cell Discov. 2023;9(1):101.

65. Hambrook JR, Hanington PC. Immune Evasion Strategies of Schistosomes. Front Immunol. 2020;11:624178.

66. McIntyre TM, Prescott SM, Stafforini DM. The emerging roles of PAF acetylhydrolase. J Lipid Res. 2009;50 Suppl(Suppl):S255–9.

67. Schilke RM, Blackburn CMR, Bamgbose TT, Woolard MD. Interface of Phospholipase Activity, Immune Cell Function, and Atherosclerosis. Biomolecules. 2020;10(10).

68. Stremler KE, Stafforini DM, Prescott SM, McIntyre TM. Human plasma platelet-activating factor acetylhydrolase. Oxidatively fragmented phospholipids as substrates. J Biol Chem. 1991;266(17):11095–103.

69. Koenig W, Khuseyinova N, Löwel H, Trischler G, Meisinger C. Lipoprotein-associated phospholipase A2 adds to risk prediction of incident coronary events by C-reactive protein in apparently healthy middle-aged men from the general population: results from the 14-year follow-up of a large cohort from southern Germany. Circulation. 2004;110(14):1903–8.

70. Stafforini DM, Carter ME, Zimmerman GA, McIntyre TM, Prescott SM. Lipoproteins alter the catalytic behavior of the platelet-activating factor acetylhydrolase in human plasma. Proc Natl Acad Sci U S A. 1989;86(7):2393–7.

71. Krieg R, Jortzik E, Goetz AA, Blandin S, Wittlin S, Elhabiri M, et al. Arylmethylamino steroids as antiparasitic agents. Nat Commun. 2017;8:14478.

72. Golan DE, Brown CS, Cianci CM, Furlong ST, Caulfield JP. Schistosomula of Schistosoma mansoni use lysophosphatidylcholine to lyse adherent human red blood cells and immobilize red cell membrane components. J Cell Biol. 1986;103(3):819–28.

73. van der Kleij D, Latz E, Brouwers JF, Kruize YC, Schmitz M, Kurt-Jones EA, et al. A novel host-parasite lipid cross-talk. Schistosomal lyso-phosphatidylserine activates toll-like receptor 2 and affects immune polarization. J Biol Chem. 2002;277(50):48122–9.

74. Gaschler MM, Stockwell BR. Lipid peroxidation in cell death. Biochem Biophys Res Commun. 2017;482(3):419–25.

75. Bennett M, Gilroy DW. Lipid Mediators in Inflammation. Microbiol Spectr. 2016;4(6).

76. Vial HJ, Torpier G, Ancelin ML, Capron A. Renewal of the membrane complex of Schistosoma mansoni is closely associated with lipid metabolism. Mol Biochem Parasitol. 1985;17(2):203–18.

77. Liu R, Cheng WJ, Tang HB, Zhong QP, Ming ZP, Dong HF. Comparative Metabonomic Investigations of From SCID Mice and BALB/c Mice: Clues to Developmental Abnormality of Schistosome in the Immunodeficient Host. Front Microbiol. 2019;10.

78. Pereira AS, Padilha RJ, Lima-Filho JL, Chaves ME. Scanning electron microscopy of the human low-density lipoprotein interaction with the tegument of Schistosoma mansoni. Parasitol Res. 2011;109(5):1395–402.

79. Huang SC, Freitas TC, Amiel E, Everts B, Pearce EL, Lok JB, et al. Fatty acid oxidation is essential for egg production by the parasitic flatworm Schistosoma mansoni. PLoS Pathog. 2012;8(10):e1002996.

80. Kadesch P, Quack T, Gerbig S, Grevelding CG, Spengler B. Tissue- and sex-specific lipidomic analysis of Schistosoma mansoni using high-resolution atmospheric pressure scanning microprobe matrix-assisted laser desorption/ionization mass spectrometry imaging. PLoS Negl Trop Dis. 2020;14(5):e0008145.

